# Oyster: A Neural Network for Modeling Genomic Sequences with Exact Position-Specific *k*-mer Contributions

**DOI:** 10.1101/2025.11.01.686014

**Authors:** Husam Abdulnabi, J. Timothy Westwood

**Author notes:** Corresponding author (HA).

## Abstract

Genomics data are sequences, and exploration in these data involves mapping these sequences to function and understanding that mapping. Analogous to a sentence composed of words, a sequence’s function is understood through its constituent subsequences, or *k*-mers. The presence of *k*-mers, their positions, and their relationships with other *k*-mers (*k*-mer associations) all affect sequence function. Understanding the sequence function requires determining the contributions of each of its *k*-mers and the impacts of their positions and *k*-mer associations.

*k*-mer associations are inherently combinatorial and may be hierarchical. Although these associations may be impactful for biological phenomena, *k*-mer associations cannot be understood by humans and severely obfuscate model analyses, yielding generalizations, simplifications, and approximations. Ignoring *k*-mer associations, however, yields concepts that are human-understandable and exact. To determine whether ignoring *k*-mer associations is appropriate, models that learn or ignore *k*-mer associations can be compared.

Oyster is a novel NN that learns *k*-mer composition and *k*-mer positions while ignoring *k*-mer associations, the first with these abilities to our knowledge. Oyster is a glass-box model as the exact position-specific contributions of *k*-mers are directly extracted from the model weights.

Oyster is compared with models that consider *k*-mer associations using two case studies spanning two important genomics topics: protein-DNA interactions and enhancer activity. Across these case studies it was found that while models that consider *k*-mer associations perform better than Oyster models, the predictions of the Oyster models were in strong agreement, serving as faithful proxies. Human-understandable concepts were extracted from the Oyster models revealing concepts that align with the literature as well as novel ones.

## 1 Introduction

Phenomena in genomics may be too complex to be fully understood by humans, thereby requiring simplifications. Although genomic phenomena can be effectively modeled through Machine learning, the models traditionally used are typically too complex to be readily interpreted. Simpler models limited to human-understandable concepts may perform sufficiently well compared to complex models, ultimately providing greater utility.

The requirements and challenges involved in understanding phenomena in genomics and the necessary simplifications for human understanding are explored in **Section 1.1**. **Section 1.2** reviews current Machine learning methods and identifies the absence of a model that can isolate the importance of subsequences and their locations from the effects of their complex interactions. To address this gap, a novel Neural Network named Oyster is introduced in **Section 2**, which is designed to model these fundamental components while explicitly ignoring their associations. Oyster is demonstrated by modelling protein-genome interactions, specifically those of the CTCF transcription factor in **Section 3** and modelling enhancer activity in **Section 4**. Human-understandable concepts are extracted from an Oyster model revealing concepts that align with current literature as well as novel ones.

### 1.1 Understanding Phenomena in Genomics

This study defines genomics as the study encompassing nucleotide (nt) sequences (DNA and RNA) and the molecules that interact with them. In addition to nt sequences, genomics data includes scores or attributes that correspond to nts of a reference nt sequence, referred to as signals (**Fig. 1**), which typically derive from physical experiments such as Chromatin Immunoprecipitation. As signals map to nt sequences, they are sequences themselves. Nt sequences and signals are hereafter collectively referred to as sequences.

**Fig. 1.**
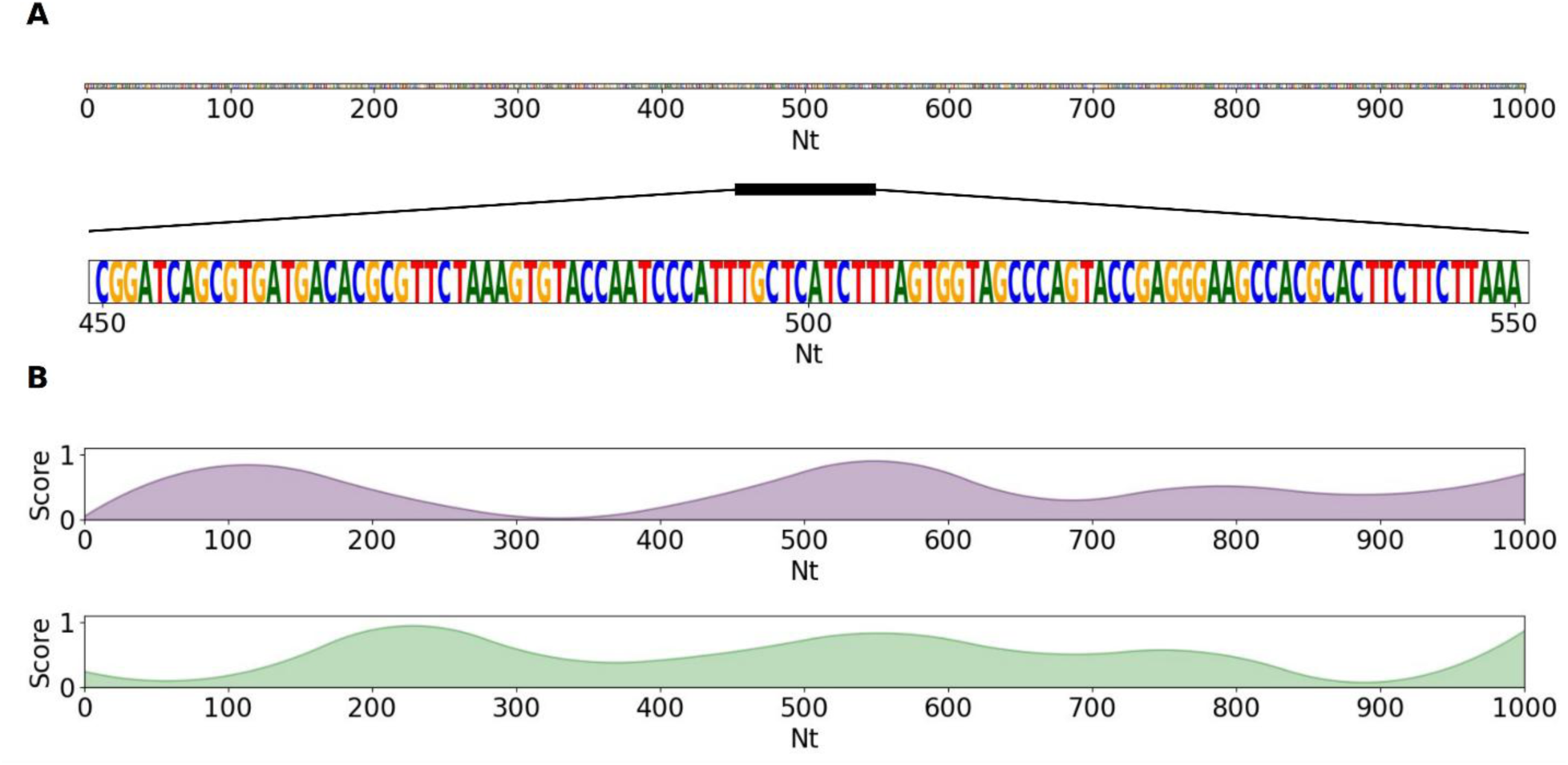
Genomics data includes nucleotide sequences and signals that correspond to them. **A)** Example of a 1000 nucleotide (nt) sequence. The center 100 nts are expanded to display the nts. **B)** Examples of two signals that correspond to the nt sequence in **A**: signal 1 (top, purple) and signal 2 (bottom, green).

Understanding genomic data involves mapping sequences to signals that correspond to the same nt(s) of a reference nt sequence. Sequences can be hundreds of millions of nts long (e.g. human chromosome 1 is 250 million nts long). Sequences are often fragmented into shorter overlapping sequences (i.e. windows) which can vary in size for each sequence used. Windows of a signal are mapped as a function of corresponding windows (i.e., windows centered on the same reference nt) of other sequences (**Fig. 2**). A function of a sequence in genomics derives from the overlapping subsequences of the sequence, analogous to a function of a sentence deriving from the words of the sentence in natural language. For simplicity, subsequences of a sequence are of the same length *k*, referred to here as *k*-mers. Hence, a function of a sequence is a function of its *k*-mers.

**Fig. 2.**
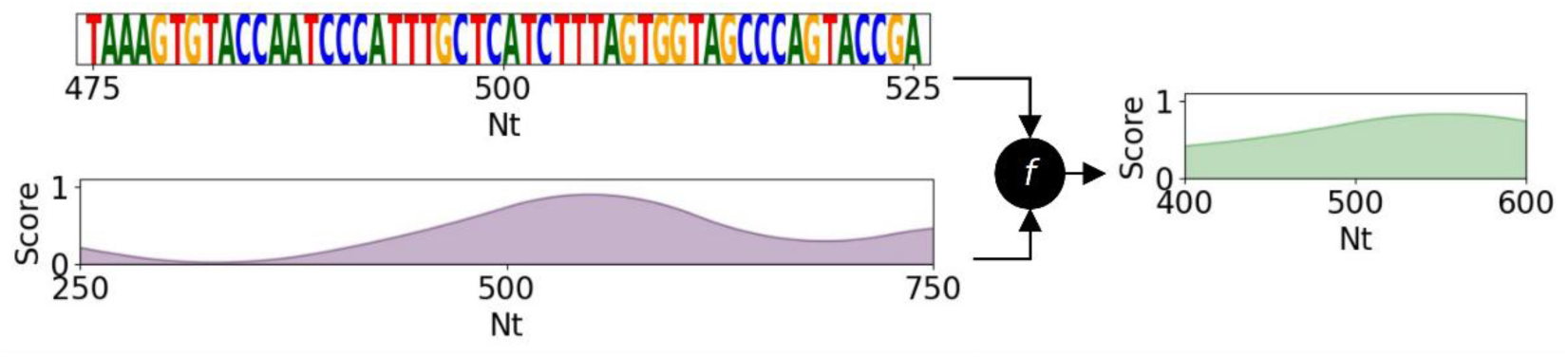
A window of a signal is a function of corresponding windows of another sequence(s). Following the example of Fig. 1, a 200 nt window of signal 2 is a function (*f*) of a corresponding 50 nt window of the nt sequence and a corresponding 500 nt window of signal 2. The differing window lengths simply illustrate that various window sizes can be used.

*k*-mers vary in their roles primarily by their interactions with other molecules (e.g. nt *k*-mers interacting with proteins). Accordingly, *k*-mers are expected to vary in their contribution to a function of a sequence. Along with their composition, the contribution of a *k*-mer may depend on its position in the sequence and/or its interactions with other *k*-mers in the sequence. Due to the nature of modelling windows of a reference sequence, the positions of *k*-mers are often crucial. The interactions among *k*-mers, referred to here as *k*-mer associations, are naturally combinatorial and can be hierarchical (Chen et al., 2014).

Understanding the mapping of sequence(s) to function requires determining the contribution of each *k*-mer of the sequence(s). It also requires determining the impacts of the *k*-mer position and *k*-mer associations on the contribution of each *k*-mer. This is accomplished at two scales: local and global.

The local scale examines each individual instance of a sequence(s)-to-function in isolation. When considering *k*-mer associations, understanding the contributions of *k*-mers is difficult even with short sequences. With as few as 10 *k*-mers and discounting potential hierarchies, there are over 1,000 potential combinations of *k*-mers to consider (2^10^ − 1). In reality, sequences in genomics are typically hundreds if not thousands of nts long, composed of hundreds if not thousands of *k*-mers. Consequently, simplification is required: instead of considering all *k*-mers and their associations, only the pairwise *k*-mer associations of the most contributive *k*-mers are explored (e.g. de Almeida et al., 2022).

The global scale seeks to derive universal relationships of all possible *k*-mers and their interactions in all possible sequences. At this scale, *k*-mer associations must consider *k*-mer positions. For a relationship of *D k*-mers, the impact of a single *k*-mer association on the contribution of a single *k*-mer is dependent on the positions of all *k*-mers involved. This is mapped on an *D* + 1 dimension space where the dimensions are the positions of each *k*-mer and the last dimension is the impact of *k*-mer association on the contribution of a single *k*-mer. Since each *k*-mer is dependent on all other *k*-mers in the sequence, this information is derived from determining the impact of a specific *k*-mer association for every possible combination of positions in every possible background sequence and summarizing them, ultimately yielding a generalization.

Besides being a generalization, even tracking position-dependent pairwise relationships of *k*-mers (again discounting potential hierarchies) scales at O(*M*^2^), where *M* is the number of *k*-mers. While one of these relationships is human-understandable, the sheer number of these relationships surpasses human understanding. In practice, the position-dependent pairwise relationships of only the most influential *k*-mers are tracked, which is far from sufficiently exhaustive of all pairwise relationships, let alone the other combinations of *k*-mers and considering hierarchies.

Until now, it was assumed that the information for the contribution of a *k*-mer and the impacts of *k*-mer position and *k*-mer associations are reasonably attainable and accurate. However, in practice, they are difficult to obtain and are essentially estimations (Ancona et al., 2019; Koo et al., 2021). Regardless, *k*-mer associations are the principal obstacle to understand genomics phenomena, requiring drastic simplifications and involving generalizations and approximations.

Ignoring *k*-mer associations—that is, only considering *k*-mer composition and *k*-mer position—immensely facilitates human understanding of genomics phenomena. Since a *k*-mer contribution is only dependent on its position, a global scale understanding can be immediately determined as the contribution of a *k*-mer at each position. Since it is not dependent on any other *k*-mer, the contributions of a *k*-mer across all possible positions are single exact values (i.e. not an approximation). This can be determined for all possible *k*-mers and be well within the bounds of human comprehension.

It is not known *a priori* whether ignoring *k*-mer associations is appropriate. Machine learning provides an opportunity to determine whether ignoring *k*-mer associations is justifiable by comparing the performance of models that consider or ignore *k*-mer associations.

### 1.2 Machine Learning Genomic Sequences

Machine learning (ML) models, especially Neural Networks (NNs), have been effective in modelling genomic sequences. All ML approaches for modeling genomic sequences can be distilled into three fundamental steps: *k*-mer embedding, sequence embedding, and output mapping (explored in Abdulnabi & Westwood, 2026).

*k*-mer embedding projects a *k*-mer as a *v*-dimensional vector where similar *k*-mers occupy similar space. *k*-mer embedding can be function-agnostic or function-specific. Function-agnostic *k*-mer embedding is performed outside the model. There are three common methods for function-agnostic *k*-mer embeddings: gapped *k*-mer counts (Ghandi et al., 2014), Transcription Factor (TF) motif counts/scores (Beer & Tavazoie, 2004), and dna2vec (Ng, 2017). Function-specific *k*-mer embeddings are learned end-to-end within the model and are typically more expressive than function-agnostic embeddings. This is best accomplished by NNs, specifically using convolutional layers. *k*-mer embedding of a sequence yields a (*L* − *k* + 1, *v*) matrix, where *L* is the length of the sequence.

Sequence embedding summarizes the *k*-mer embeddings to a *q*-dimensional vector that can take into consideration *k*-mer position and *k*-mer association. Sequence embedding can be function-agnostic or function-specific. Function-agnostic sequence embedding involves summarizing the sequence with non-learnable methods. One common method is by summing the *k*-mer embeddings by embedding dimension which ignores *k*-mer position and *k*-mer associations. Function-specific sequence embedding involves learnable methods. There are several different methods for function-specific sequence embedding (described in Abdulnabi & Westwood, 2026) which differ in their capacity to capture *k*-mer associations and positional dependencies.

Output mapping takes the sequence embedding and maps it to an output and is always within the model. Output mapping can ignore further *k*-mer associations by a linear combination of the sequence embedding vector (i.e., weighted sum) followed by a link function which can be linear or non-linear.

To our knowledge, no formalized ML method currently ignores *k*-mer associations while considering *k*-mer positions. Consequently, there remains a gap for a framework that combines these constraints with the power of function-specific *k*-mer embeddings.

To address this, we developed Oyster: the first NN that ignores *k*-mer associations while considering *k*-mer positions and performs function-specific *k*-mer embedding. Position-specific *k*-mer contributions are extracted directly and easily from the model. We compare Oyster to powerful models that consider *k*-mer associations in two areas of genomics research: protein-DNA interactions and enhancer activity. We find that Oyster predictions align with those of the other models, justifying its use. We then determine the position-specific *k*-mer contributions, revealing concepts that align with established literature while revealing novel regulatory rules.

## 2 Oyster

Oyster is a novel NN for modelling genomic sequences that ignores *k*-mer associations while considering *k*-mer positions and performs function-specific embedding. Oyster is named after the pearls of exact human-understandable concepts that can be extracted from them.

The position-specific contributions of a given *k*-mer can be easily determined directly from an Oyster architecture, obviating the need for post-hoc attribution methods that are typically required for established black-box NNs. Thus, Oyster is a glass-box model. Moreover, while most established NNs can only handle nt sequences, Oyster can handle both sequences and signals and can also handle multiple sequences simultaneously.

The Oyster architecture is explored in **Section 2.1**. A single Oyster can handle a nt sequence or multiple signals with the same length. To model both nt sequence and signal(s) or signals with different lengths, multiple Oysters can be used and combined in a single architecture called Multi-Oyster, explored in **Section 2.2**. The analysis of Oyster is explored in **Section 2.3**. Oyster software and a manual with more details are provided in https://github.com/Husam94/poseigen_oyster/.

### 2.1 Architecture

The Oyster architecture (**Fig. 3**) has 3 main steps: *k*-mer embedding (**Section 2.1.1**), positional reduction (**Section 2.1.2**), and output (**Section 2.1.3**). The Oyster architecture derives from the Mimic architecture (Abdulnabi & Westwood, 2026b).

**Fig. 3.**
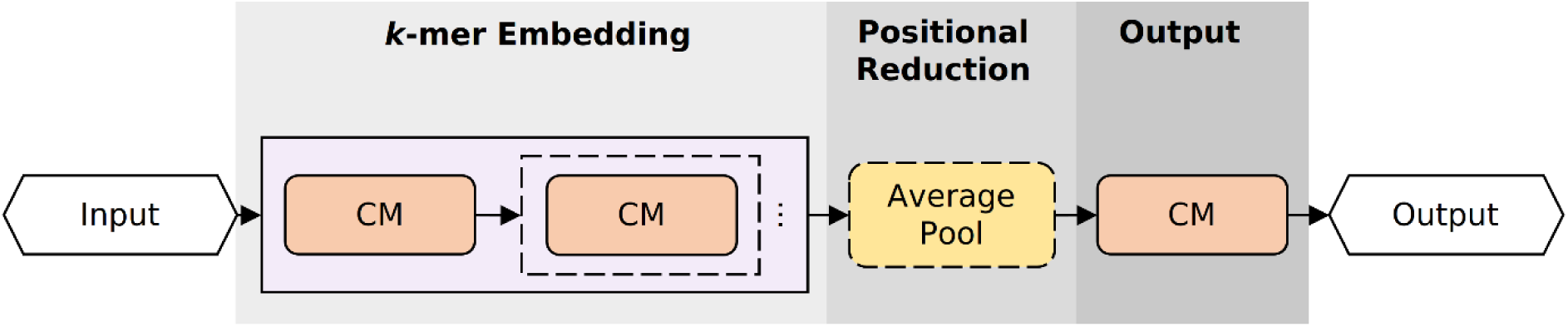
Oyster architecture. An input is a sequence oriented 5’-3’. *k*-mers of the sequence(s) are embedded using 1 or more convolutional modules (CMs) that are composed of a convolutional layer followed by an optional activation function. The length of the *k*-mer embeddings is optionally reduced using non-overlapping average pooling. The reduced *k*-mer embeddings are then mapped to the output by 1 CM. Boxes with dashed lines are optional operations and 3 dots indicate repeated operations.

#### 2.1.1 *k*-mer Embedding

Oyster accepts either nt sequences or signals as inputs. The Oyster architecture uses two-dimensional convolutional layers to perform operations similar to one-dimensional convolutional layers which are present in other architectures (e.g. Novakovsky et al., 2023). The two-dimensional convolutional layers allot greater flexibility and functionality for modelling nt sequences or signals, as will become evident later in this section. For simplicity, the shapes of the arrays are (depth, length, width) where depth is the number of filters (i.e. channels), length is the sequence positions, and width is the features per position. The two-dimensional convolutional layers have filters that span the entire width of the incoming sequences.

Nt sequences are one-hot encoded with each nt dimension as the width which is an array of shape (1, length(*x*), 4). Multiple signals of the same length can be modelled by a single Oyster. Signals are arranged such that each signal is its own depth, which is an array of shape (*N*, length(*x*), 1).

Oyster first embeds overlapping *k*-mers of a sequence using convolutional modules (CMs) that are composed of a convolutional layer followed by an activation function. A single CM can be used with filters of length *k*. This method is similar to scoring *k*-mers by Position Weight Matrices (PWMs) and assumes each element of the *k*-mer to be independent.

To consider higher-order intra-*k*-mer dependencies, multiple CMs are used (Abdulnabi & Westwood, 2026b). Each convolutional layer uses a base filter length *m* < *k*, with the final layer using a shorter filter of length *n* < *m*. For example, to embed *k*-mers of length 14 with a base filter of 6, convolutional layers with filter lengths of 6, 6, and 4 are used. The base filter length is specified and the number of CMs to use and the filter length per CM are automatically determined internally.

The number of filters used in the last CM is specified by the number of embedding dimensions desired (*v*). The number of filters of the intermediate CMs (if applicable) are imputed by linear number spacing. *k*-mer embedding yields an array of shape (*v*, *L* − *k* + 1, 1) where L is the length of the sequence.

For a sequence composed of multiple signals, the *k*-mers of each signal can be embedded separately from other signals by using convolutional layers with grouped filters. This facilitates distinguishing the contribution of *k*-mers of each signal and the contribution of each signal in later analysis.

The size of *k*, the base filter length, the number of embedding dimensions, and whether to use grouped filters are tunable parameters (i.e. hyperparameters).

#### 2.1.2 Positional Reduction

The positional resolution of *k*-mer embeddings can be reduced by average pooling which does not consider *k*-mer associations. Reducing positional resolution can provide greater model generalizability by regularizing *k*-mer embeddings.

The desired resolution (*r*) is achieved through non-overlapping pooling which automatically determines the pooling length and stride to reduce the length to 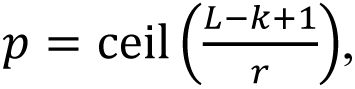 referred to as positional blocks. To facilitate analysis, the divisor for average pooling is *r* for all positional blocks. The entire length of the *k*-mer embeddings can be pooled which ignores *k*-mer position. The desired resolution is a hyperparameter. The positional reduction step yields arrays of shape (*v*, *p*, 1).

#### 2.1.3 Output

The outputs of NNs for genomic sequences are of shape (*out_lengt_*_ℎ_, *out_type_*) where *out_type_* refers to different signals. While NNs can produce outputs with *out_lengt_*_ℎ_ greater than 1, Oyster outputs have a length of 1 to facilitate later analysis (Abdulnabi & Westwood, 2026b). Oyster maps the positionally reduced *k*-mer embeddings to the output through a single CM with a convolutional layer with *out_type_* number of filters of size of (*v*, *p*, 1). Each filter is effectively a learnable weighted sum of the positionally reduced *k*-mer embeddings. This is the fundamental step in learning *k*-mer positions while ignoring *k*-mer associations.

### 2.2 Multi-Oyster

Multiple Oysters can be used in a single architecture for modelling multiple different input sequences and is called Multi-Oyster (**Fig. 4**). Each Oyster can attend to 1 or more sequences and has their own hyperparameters. For example, one Oyster attends to nt sequences and the other attends to signals. The individual Oyster models do not have a final activation function to facilitate later analysis. The outputs of the Oysters are combined additively (i.e. summed), and an optional final activation function can be applied.

**Fig. 4.**
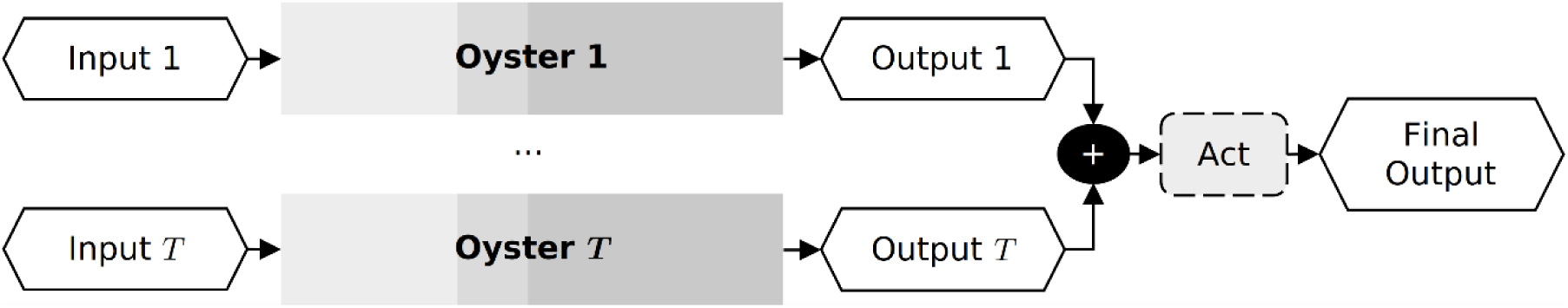
Multi-Oyster Architecture. Each input is fed through a separate Oyster model that produces a corresponding output. The outputs are summed followed by an optional activation function (“Act”). See Fig. 3 for additional details.

### 2.3 Model Analysis

The contribution of a *k*-mer to an *out_type_* at each positional block is simply the dot product of its embedding and the corresponding weights of the output convolutional layer scaled by the positional resolution (**Eq. 1**).

**Eq. 1. Position-specific *k*-mer contributions**

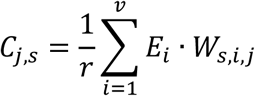

Where:

*C_j_*_,*s*_ = *k*-mer contribution at positional block *j* (*j* ∈ {1,…,*p*}) for output signal *s*

*r* = positional resolution

*E_i_* = *k*-mer embedding value for dimension *i* of *v* embedding dimensions

*W_s_*_,*i*,*j*_ = learnable weight of the output signal *s* and embedding dimension *i* at position *j*

When using multiple sequences where each sequence is handled by a separate Oyster or by grouped convolutional filters, the contribution of a sequence to an *out_type_* is the dot product of its *k*-mer embeddings and the corresponding weights of the output convolutional layer scaled by the positional resolution (**Eq. 2**).

**Eq. 2. Sequence contribution**

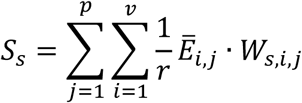

Where:

*S_s_* = total contribution of the sequence to output signal *s*

*Ē_i,j_* = sum of the embeddings within positional block *j* for dimension *i* before scaling by *r*

The contributions of a sequence can be positive or negative. The *influence* of a sequence to the output is determined as the proportion of its absolute contribution to the sum of all absolute sequence contributions (**Eq. 3**).

**Eq. 3. Sequence influence**

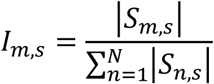

Where:

*I_m_*_,*s*_ = the influence of sequence *m* on output signal *s*

*S_m_*_,*s*_ = the total contribution of sequence *m* to output *s* (derived from **Eq. 2**)

*N* = the total number of sequences

#### 2.3.1.1 *k*- and *z*-Patterns

Determining and storing the position-specific contributions of all *k*-mers in the data (let alone all possible *k*-mers) individually may require resources (computational and time) that exceed resource limits. The *k*-mers can be reduced to clusters determined by their composition. Nt *k*-mers are represented by their one-hot encoding. Each cluster can be represented by the cluster centroid, which is the average of its members, referred to as a *k*-pattern. Notably, nt *k*-patterns are Position Probability Matrices (PPMs): probabilities of each type of nt (A, C, G, T/U) at each element.

The position-specific contribution of a *k*-pattern is determined by the position-specific contributions of its constituent *k*-mers. The contribution of a *k*-pattern at a given position is determined as the weighted average of the contributions of its constituent *k*-mers, weighted by their similarity to the *k*-pattern by the reciprocal (i.e. multiplicative inverse) of the Euclidean distance.

There may be existing patterns before modelling such as protein-DNA motifs which typically range in size from 6–18 elements. As Oyster models characterize *k*-mers of a single *k*, patterns that are not of size *k*, referred to as *z*-patterns, are modified to produce *k*-patterns, exemplified in **Fig. 5** with the nucleotide motif of the transcription factor MYC. For a *z*-pattern of size *z* < *k*, *k*-patterns are generated by sliding the *z*-pattern from positions 1 to *k* − *z* + 1. This generates *k* − *z* + 1 *k*-patterns with *k* − *z* empty positions that are filled with equal probabilities (0.25) for each nt. For a *z*-pattern of size *z* > *k*, *k*-patterns are generated by sliding a window of size *k* from positions 1 to *z* − *k* + 1 on the *z*-pattern which generates *z* − *k* + 1 *k*-patterns.

**Fig. 5.**
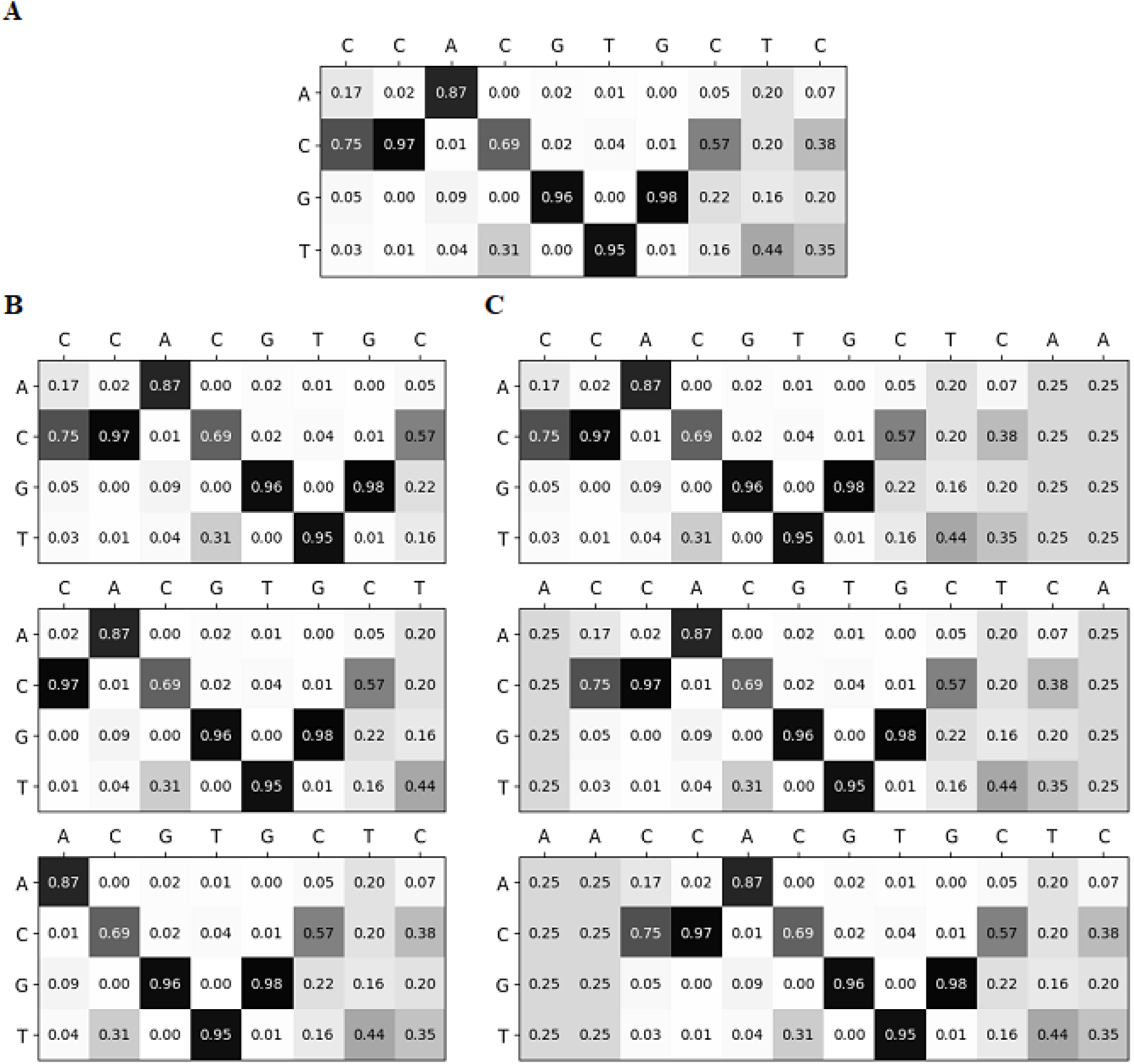
*k*-patterns of the MYC motif. **A)** MYC motif consists of 10 nts. **B)** MYC *k*-patterns with *k* of 8. **C)** MYC *k*-patterns with *k* of 12. Rows correspond to nts A, C, G, and T, respectively and values correspond to frequency. Columns for each *k*-pattern are labelled by the dominant nt. Light-dark color scale visualizes values from [0, 1].

The contribution of a *z*-pattern per position of *L* − *z* + 1 is the sum of the belonging *k*-pattern contributions at their corresponding positions. That is, the contribution of a *z*-pattern at a given position *i* is the sum of the first belonging *k*-pattern at position *i*, the second belonging *k*-pattern at position *i* + 1, etc.

## 3 Case Study 1: CTCF-Genome Interactions

The interactions of proteins to the genome, referred to as protein-genome interactions (PGIs), are central to genomics as they are the principal components of gene regulation. The frequency and strength (collectively referred to here as intensity) of an interaction of a protein to a location in the genome may depend on the underlying DNA sequence and/or the surrounding environment which includes histones and their post-translational modifications, simply referred to as histones hereafter (Guertin & Lis, 2010; Slattery et al., 2014; Wang et al., 2012).

This case study is an example of modelling PGIs of CTCF using Oyster models. CTCF is a DNA sequence-specific transcription factor influential in chromatin architecture (Phillips and Corces, 2009). The PGIs of CTCF in human cancer cells are modelled as functions of human genome DNA sequence and 11 histone-genome signals.

CTCF-genome interactions were previously modelled using three models in Abdulnabi & Westwood (2026b): TF-Lasso, Bichrom, and Multi-Mimic. TF-Lasso serves as a popular non-NN comparator (de Almeida et al., 2022). Bichrom serves as a popular NN comparator (Srivastava et al., 2021). Multi-Mimic is an NN that facilitates optimization and module comparison. Bichrom and Multi-Mimic consider *k*-mer associations. It was found that the Multi-Mimic significantly outperforms the other models and is used as the comparator in this study.

Here, CTCF-genome interactions are modelled with Multi-Oyster composed of two Oysters, one that attends to DNA sequence and the other that attends to histone signals (**Fig. 6**). Multi-Oyster is compared to the Multi-Mimic model. Model analysis is conducted on the Multi-Oyster model.

**Fig. 6.**
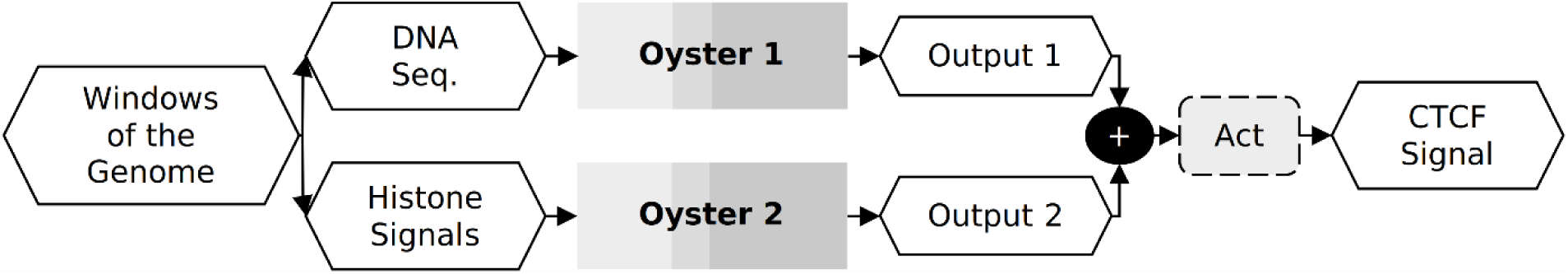
CTCF signal of a window of the genome is predicted by a Multi-Oyster that uses DNA sequence and histone signals of corresponding windows of the genome. The Multi-Oyster is composed of two Oysters, one attending to DNA sequence and the other attending to histone signal. The Oyster outputs are combined to produce the final prediction of CTCF signal.

The workflow for CTCF-genome interactions was also implemented for YY1-genome interactions, yielding comparable results. For brevity and to avoid redundancy, the analysis of YY1-genome interactions is not included here but the analysis is accessible for further review at https://github.com/Husam94/CS-YY1/.

### 3.1 Methods

See https://github.com/Husam94/CS-CTCF/ for full code. See Abdulnabi & Westwood (2026b) for additional descriptions. Only relevant information is included here.

#### 3.1.1 Data Processing

The CTCF-genome and 11 histone-genome interactions were determined by Chromatin Immunoprecipitation (ChIP) followed by high-throughput sequencing. The data were retrieved from the ENCODE database (ENCODE Project Consortium, 2012).

Human genomic DNA sequences of 500 nts and histone signals of 2000 nts at a 50 nt resolution (signal of length 40) were used. Of the possible locations (i.e. sites) in the genome, 20,000 were selected based on specific criteria and sampling. Neighboring windows of ±150 nts were added, totalling 60,000 observations.

The CTCF-genome intensities are right-skewed. The density across the range of CTCF-genome interactions were determined through binning-based methods (Abdulnabi & Westwood, 2026a).

The observations were divided into training, validation, testing, and an additional hold-out set for final evaluation, with proportions of 0.55, 0.15, 0,15, and 0.15, respectively.

#### 3.1.2 Model Fitting and Scoring

Following the optimization procedure in Abdulnabi & Westwood (2026a), Multi-Oyster was optimized by a two-step procedure. The first step is random search with 300 candidates. The second step retrieves the top 10 candidates based on the performance on the validation set and conducts 10 model repeats for each candidate. The top candidate is determined as the one with the best average score on the validation set among its repeats.

Following the scoring procedure in the Abdulnabi & Westwood (2026a), an additional 5 repeats were done for the Multi-Oyster model, totalling 15 repeats. For robustness, the repeats were bootstrapped 500 times to simulate 500 bootstrapped repeat sets. For each set of bootstrapped repeats, the top 3 models (based on the score on the validation set) are determined. Across all bootstrapped repeats, the model indices of the top models are used to determine a weighting by scaling the frequency by the total number of model indices. The predictions of the model repeats are combined by weighted average, producing ensemble predictions.

The ensemble predictions are assessed by reciprocal-density weighted Pearson Correlation Coefficient (recip-PCC). The reciprocal-density weighting accounts for the right-skewed imbalance.

### 3.2 Oyster Analysis

The weighted average ensemble of the Oyster models was used for model analysis. The contributions of sequences and *k*-mers are determined for each model of the ensemble and then weighted averaged.

#### 3.2.1.1 Contributions and Influences of Sequences

For each observation, the contributions of DNA sequence and each histone signal are determined following **Eq. 2**. For each observation, the influences of DNA sequence and each histone signal are determined following **Eq. 3**. For summarizing the influences across the range of CTCF-genome intensities, the median influence of each signal is determined for each CTCF-genome bin.

#### 3.2.1.2 Contributions of Protein-DNA Motifs and Histone *k*-patterns

The position-specific contributions of *k*- and *z*-patterns are determined following **Section 2.3.1.1.** Existing protein-DNA motifs were retrieved from the HOCOMOCO database, specifically from version 11 of the core human motifs (Kulakovskiy et al., 2018). For each motif, the contribution of it and its reverse complement were determined and averaged. The contribution of a motif was determined as the sum of the contributions of its belonging *k*-patterns. For each *k*-pattern of a motif, 10,000 *k*-mers were sampled and their position-specific contributions were determined.

For each histone signal, *k*-mers (of size *k* that was used by the model) from the data used were clustered into 100 clusters using K-means clustering. The clusters were reduced using Agglomerative clustering with a distance cut-off determined by identifying the elbow/knee point by Kneedle algorithm through the *Kneed* python package (Satopaa et al., 2011). For each cluster, member profiles were averaged to define cluster centroids.

### 3.3 Results and Discussion

Although Multi-Mimic outperforms Multi-Oyster on modelling CTCF-genome interactions (**Fig. 7**), predictions from both models align strongly on the test set with a recip-PCC of 0.935. This high concordance justifies the use of Multi-Oyster as a faithful and interpretable proxy for the Multi-Mimic model, validating subsequent model analysis. It also indicates that while CTCF-genome interactions may involve complex *k*-mer associations, these interactions exert minimal impact relative to the primary effects of *k*-mer composition and positional arrangement. Additionally, Multi-Oyster maintains robust performance across all data partitions, including the training, validation, test, and holdout sets (**Sup.Fig. 1**).

**Fig. 7.**
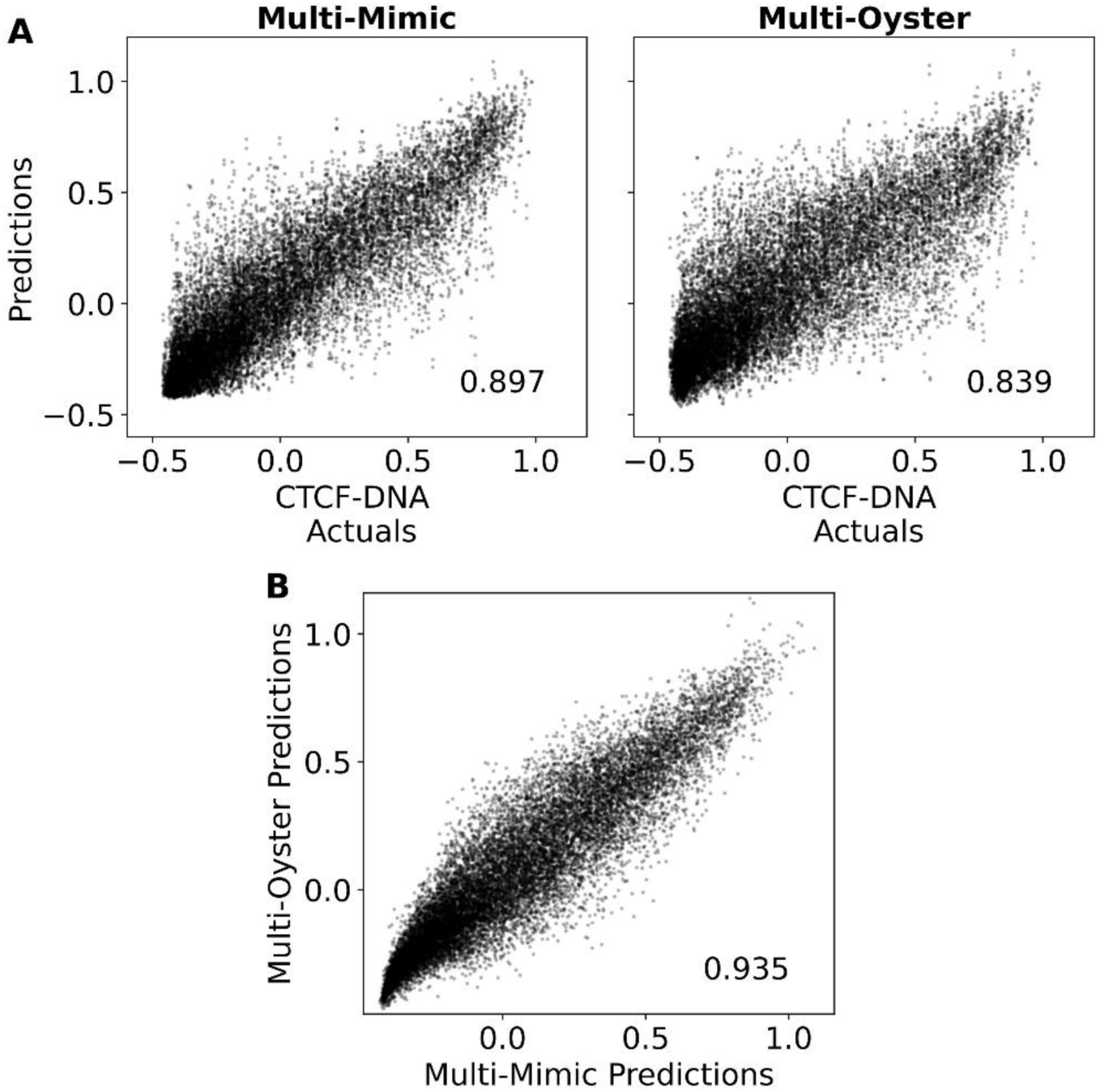
Multi-Mimic outperforms Multi-Oyster on modelling CTCF-genome interactions, but their predictions align strongly with each other. Scatter plots of weighted ensemble predictions of Multi-Mimic and Multi-Oyster on CTCF-genome interactions on the testing set compared to (**A**) actuals and (**B**) each other. The recip-PCC is included at the bottom right for each plot.

Among the sequences used to model CTCF-genome interactions, DNA sequence is dominantly influential across the entire range of actuals especially for stronger CTCF-genome interactions **(Fig. 8**). Conversely, all histones are weakly influential across the entire range of actuals, especially for stronger CTCF-genome interactions.

**Fig. 8.**
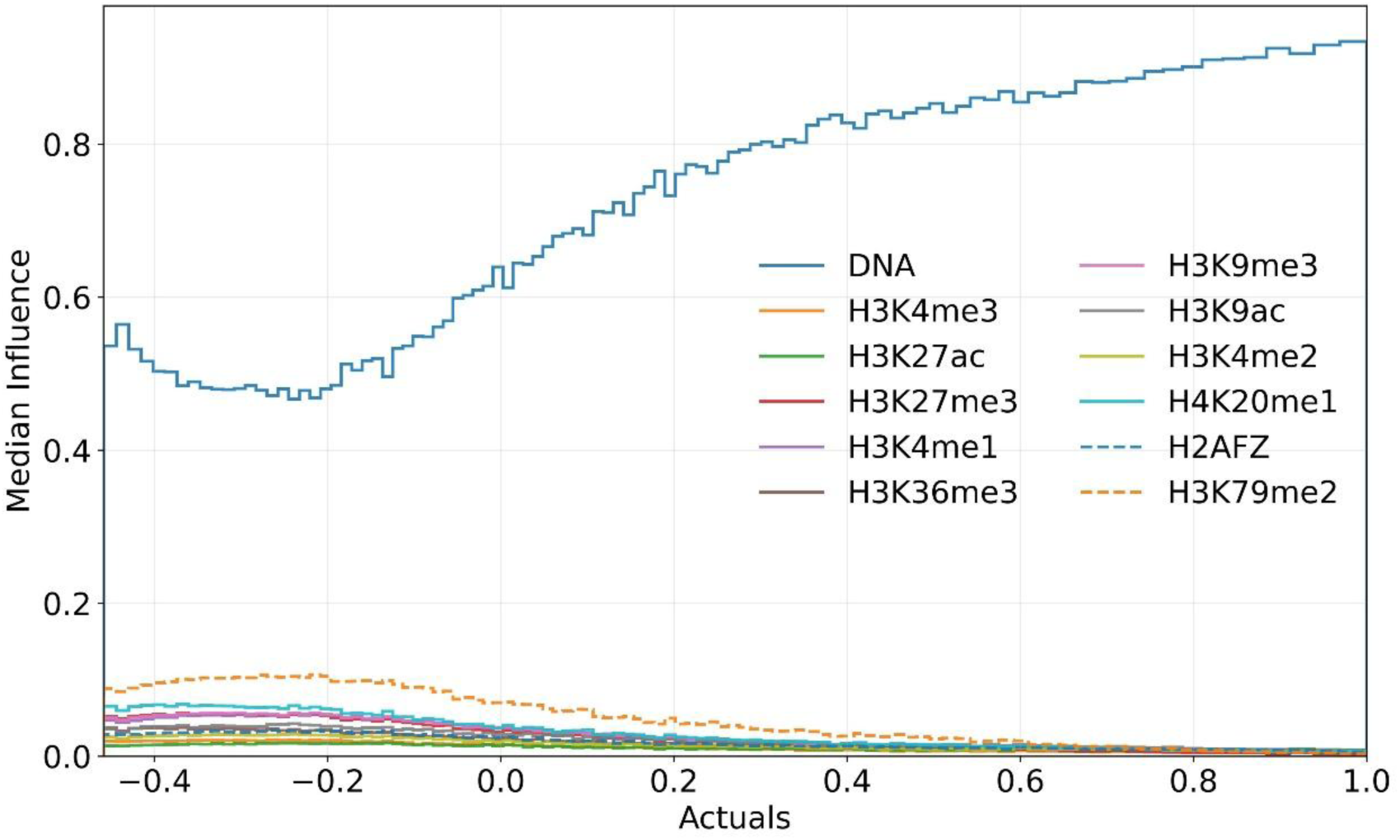
DNA is the most influential sequence for modelling CTCF-genome interactions. Stair plots visualizing the median influence of each biological sequence to the predictions pertaining to each bin across the range of actuals.

The are few strongly positive contributive motifs for modelling CTCF-genome interactions and several weakly negative contributive motifs (**Fig. 9**). The known CTCF motif is the most contributive motif, contributing positively to CTCF-genome interactions, as expected (**Fig. 10**). While this finding that CTCF motif strongly affects CTCF-genome interactions predictions is not a surprise biologically, the modelling did not have this prior, showing that Oyster recovers known biological phenomena. Thus, this finding serves as a positive control.

**Fig. 9.**
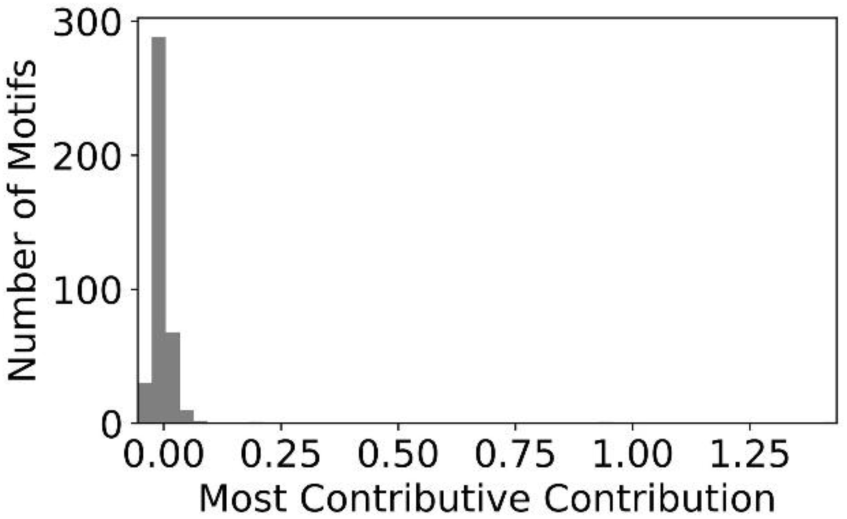
There are few strongly contributive motifs for modelling CTCF-genome interactions. Histogram with 50 bins of the most contributive contributions of the protein-DNA motifs. For each motif, the most contributive contribution is determined as the contribution at the maximum of the absolute contributions across all positions.

**Fig. 10.**
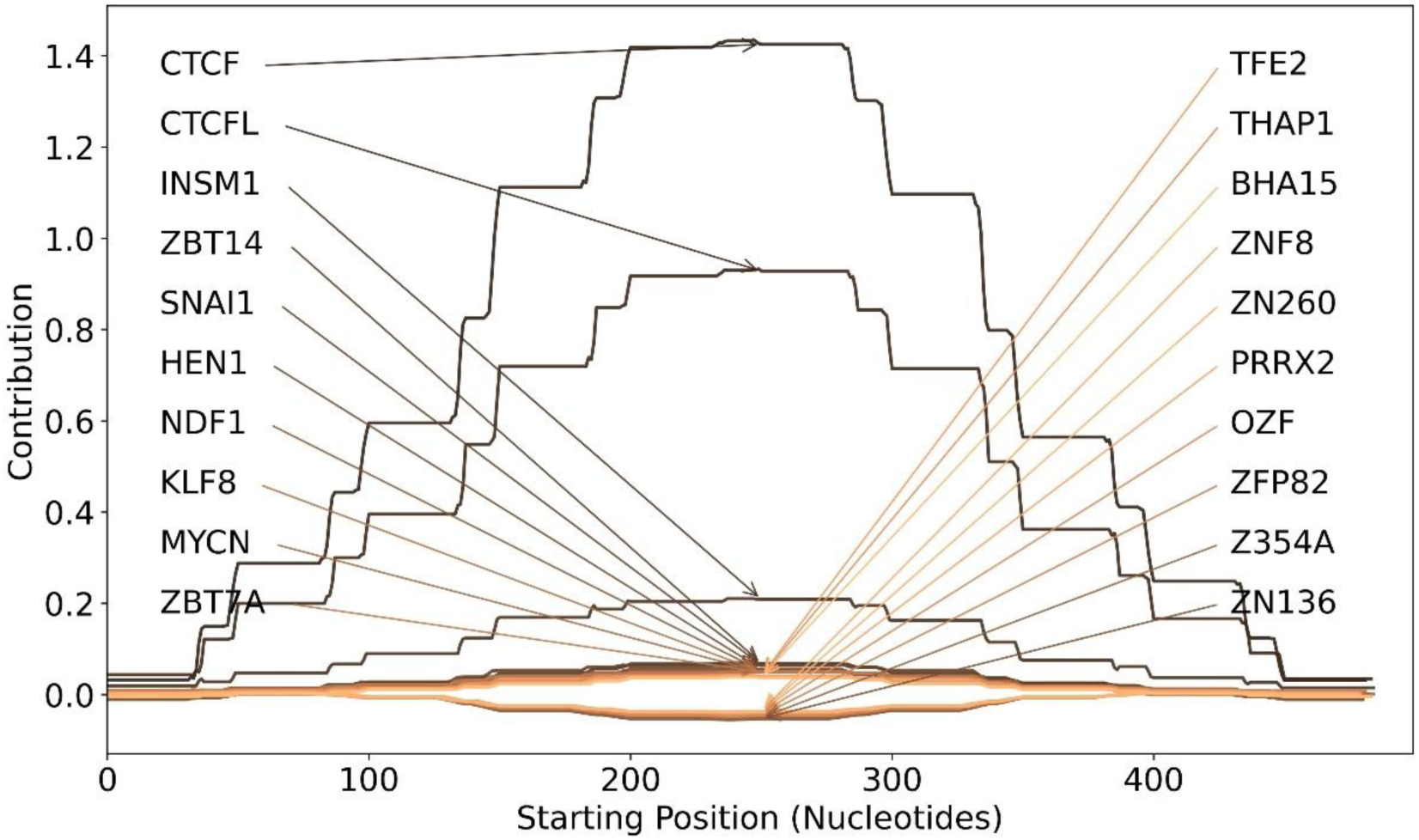
CTCF is the most contributive motif for CTCF-genome interactions. Motif contributions of the top 20 most contributive motifs for each possible starting position of the 500 nt DNA sequence.

The CTCF motifs (CTCF and CTCFL) are much more contributive than any other motif. The other substantially contributive motif is the ISM1 motif but this is likely due to its resemblance to the CTCF motif (**Fig. 11**). Interestingly, there are several negatively (albeit weakly) contributive motifs including ZN136. ZNF136 is a KRAB-Zinc Finger protein that is known to recruit mediators of H3K9me3 formation and DNA methylation that serve to block protein-DNA interactions (O’Geen et al., 2007). However, as explored earlier, H3K9me3 is minimally influential relative to DNA sequence. This warrants further investigation.

**Fig. 11.**
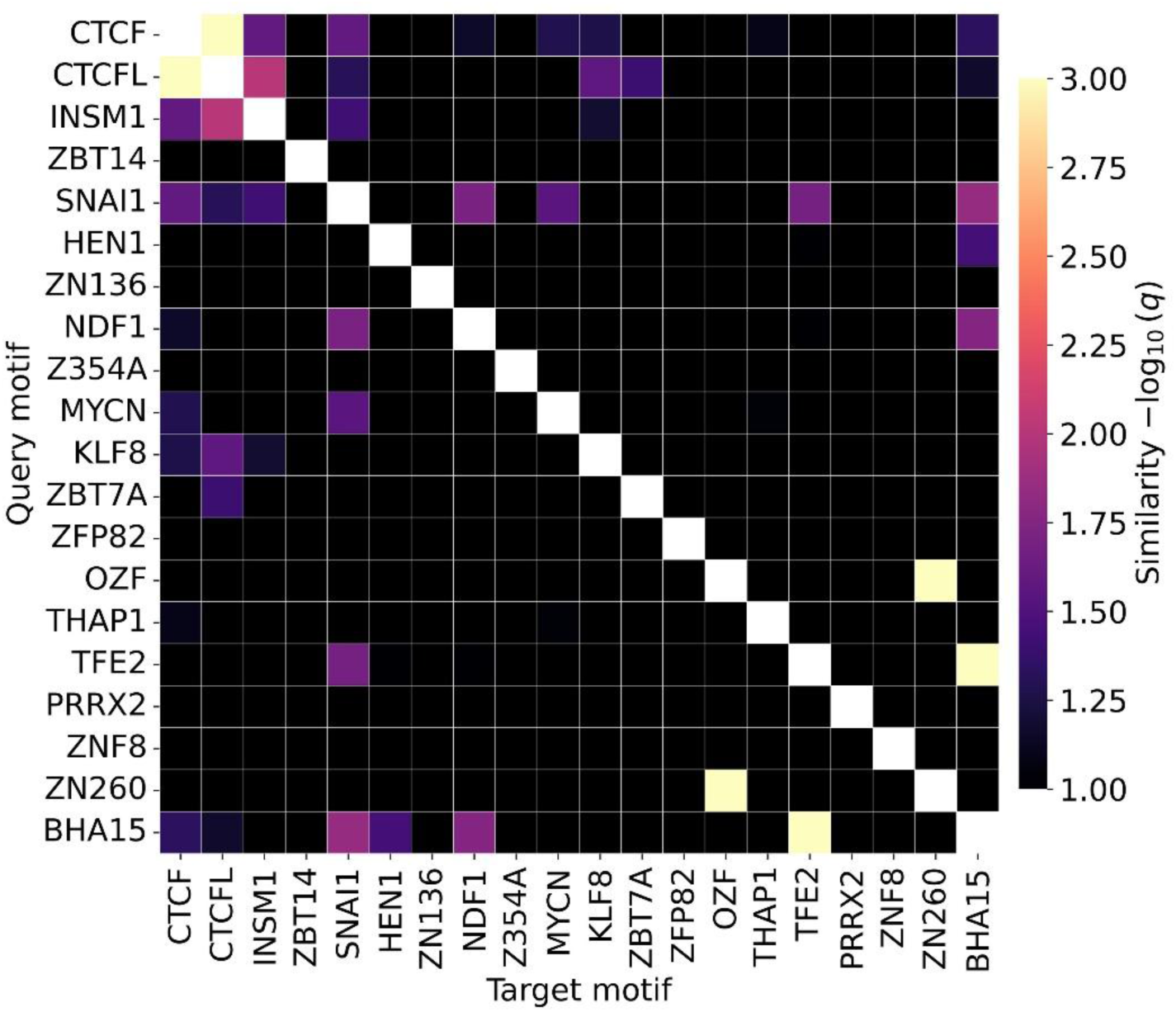
The INSM1 motif resembles the CTCF motif. TOMTOM similarity of the top 20 most impactful motifs. Dark light color scale visualizes values from 1.0 or less to 3.0 and above. A similarity of 1.0 or less is deemed non-similar.

Relative to motifs, all histone *k*-patterns are very weakly contributive (**Fig. 12**). For brevity, this study does not further explore the contributions of histone *k*-patterns but the analysis is provided in the code.

**Fig. 12.**
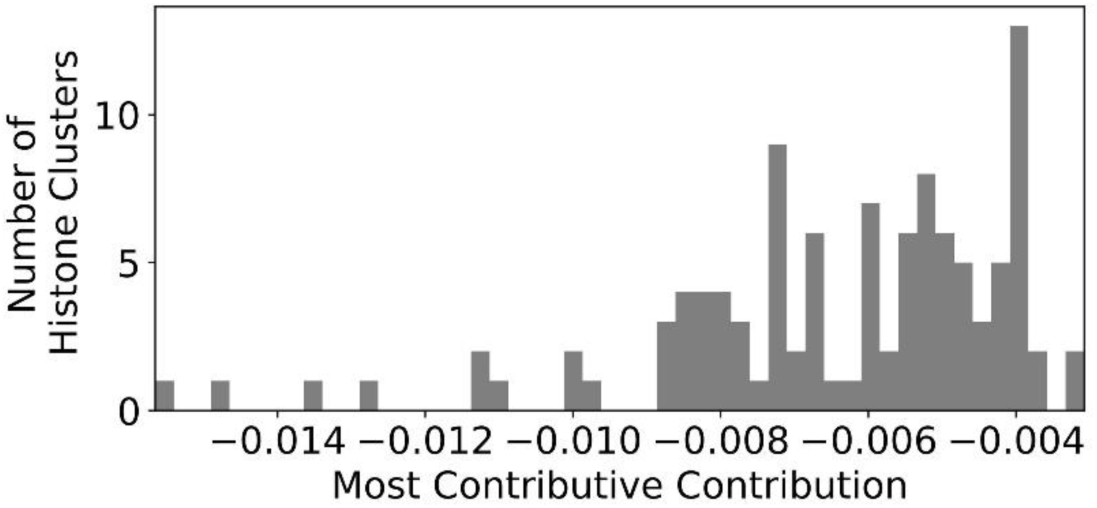
Histone *k*-patterns are weakly contributive for modelling CTCF-genome interactions. Following Fig. 9, histogram with 50 bins of the most contributive contributions of the histone *k*-patterns.

The finding that histone signals are weakly influential for CTCF-genome interactions may stem from the modeling objective and the data used. The objective is to model CTCF-genome interactions genome-wide without prior knowledge of the CTCF motif. The CTCF protein is dependent on the CTCF motif for interacting with the DNA. Relative to the entire genome, there are few regions in the genome that contain the CTCF motif.

The CTCF-genome interaction intensities genome-wide are right-skewed: there are increasingly few high intensities away from the mean, with most intensities close to 0. Due to computing resources, of the possible sites to include for modelling, 20,000 sites and their direct neighbors were selected. Site selection prioritized sites with elevated binding intensities while sampling representative sites from lower-intensity regions. Most if not all sites with higher intensities likely have CTCF motifs whereas only a relative few sites with lower intensities have CTCF motifs. Sites that harbor a CTCF motif but have low intensity are likely dependent on the chromatin environment. However, since there are likely relatively few of those sites, the model can incorrectly predict a high intensity without substantially affecting performance metrics which evaluate the predictions as an aggregate.

Moreover, the DNA sequence itself affects and therefore encodes chromatin environment information. While 2000 nt of histone signal and 500 nt of DNA sequence are used, the 500 nt sequence may sufficiently define the local chromatin environment, rendering the histone signals redundant.

One potential avenue is to model only sites that contain a CTCF motif. In this context, histones are expected to have more influence and the CTCF motif is expected to be less contributive since every site contains it. However, the resulting model would likely perform poorly genome-wide on sites devoid of a CTCF motif.

Another avenue is to incorporate the individual outputs of Oyster sub-models directly into the loss function formulation. Predictions that have higher DNA sub-model output can be weighted greater, thereby amplifying wrong predictions for sites that contain a CTCF motif but have low intensity. This forces the model to depend more on histone signals for accurate predictions.

This issue is not isolated to CTCF-genome interactions but to all DNA-sequence-specific protein-DNA interactions, especially transcription factors. Further investigations may yield very rewarding solutions that improve the understanding of the interplay between protein-DNA interactions and the chromatin environment.

## 4 Case Study 2: Enhancer Activity

Enhancer activity refers to the varying capacity of DNA sequences to regulate gene expression. A leading method for quantifying this activity is Self-Transcribing Active Regulatory Region sequencing (STARR-seq) (Arnold et al., 2013). This assay involves generating a library of plasmids containing various DNA sequences—often derived from genomic fragments— placed downstream of a minimal promoter. Upon transfection into cells, the cellular machinery transcribes these sequences; fragments with higher enhancer activity drive higher levels of their own transcription. Because this activity is measured episomally, it captures the intrinsic regulatory potential of the DNA sequence and the specific cellular state while bypassing native chromatin effects.

There are three competing theories on enhancer activity mechanisms: the Billboard, Enhancesome, and TF-collective models (Jindal & Farley, 2021). The Billboard model posits that *k*-mer associations are not required whereas the Enhancesome and TF-collective models argue that *k*-mer associations are required. Comparing Oyster models to models that consider *k*-mer associations can yield valuable insights into enhancer activity mechanisms.

This case study focuses on modelling enhancer activity in the K562 human leukemia cell line using only DNA sequence. As with the CTCF-genome interactions case study, the dataset used here was modelled before using three ML models: TF-Lasso, DeepSTARR, and Mimic. DeepSTARR is a popular NN used to model enhancer activity that is measured by STARR-seq. DeepSTARR and Mimic both consider *k*-mer associations. It was found that Mimic significantly outperforms the other models and thus is used as the comparator in this study. Here, enhancer activity is modelled using Oyster and is compared to the Mimic model. Model analysis is then conducted on the Oyster model.

### 4.1 Methods

See https://github.com/Husam94/CS-EnhAct/ for full code. See MIMIC paper for additional descriptions. Only relevant information is included here.

#### 4.1.1 Data Processing

The enhancer activity was determined using STARR-seq and the data was retrieved from the ENCODE database (ENCODE Project Consortium, 2012).

The STARR-seq used human genomic DNA sequences of roughly 800 nts. Accordingly, human genomic DNA sequences of 1000 nts were used for modelling. Of the possible locations (i.e. sites) in the genome, 30,000 were selected based on specific criteria and sampling. Neighboring windows of ±250 nts and ±500 nts were added, totalling 150,000 observations.

The enhancer activity intensities are right-skewed. The density across the range of enhancer activity intensities was determined through binning-based methods (Abdulnabi & Westwood, 2026a).

The observations were divided into training, validation, testing, and an additional hold-out set for final evaluation, with proportions of 0.55, 0.15, 0,15, and 0.15, respectively.

#### 4.1.2 Model Fitting and Scoring

Same as **Section 3.1.2** except using Oyster instead of Multi-Oyster.

#### 4.1.3 Oyster Analysis

Same as **Section 3.2.1.2** except only analyzing protein-DNA motifs.

### 4.2 Results and Discussion

Although Mimic outperforms Oyster on modelling enhancer activity (**Fig. 13**), predictions from both models align strongly on the test set with a recip-PCC of 0.948. This high concordance justifies the use of Oyster as a faithful and interpretable proxy for the Mimic model, validating subsequent model analysis. It also indicates that while enhancer activity may involve complex *k*-mer associations, these interactions exert minimal impact relative to the primary effects of *k*-mer composition and positional arrangement. Additionally, Oyster maintains robust performance across all data partitions, including the training, validation, test, and holdout sets (**Sup. Fig. 2**).

**Fig. 13.**
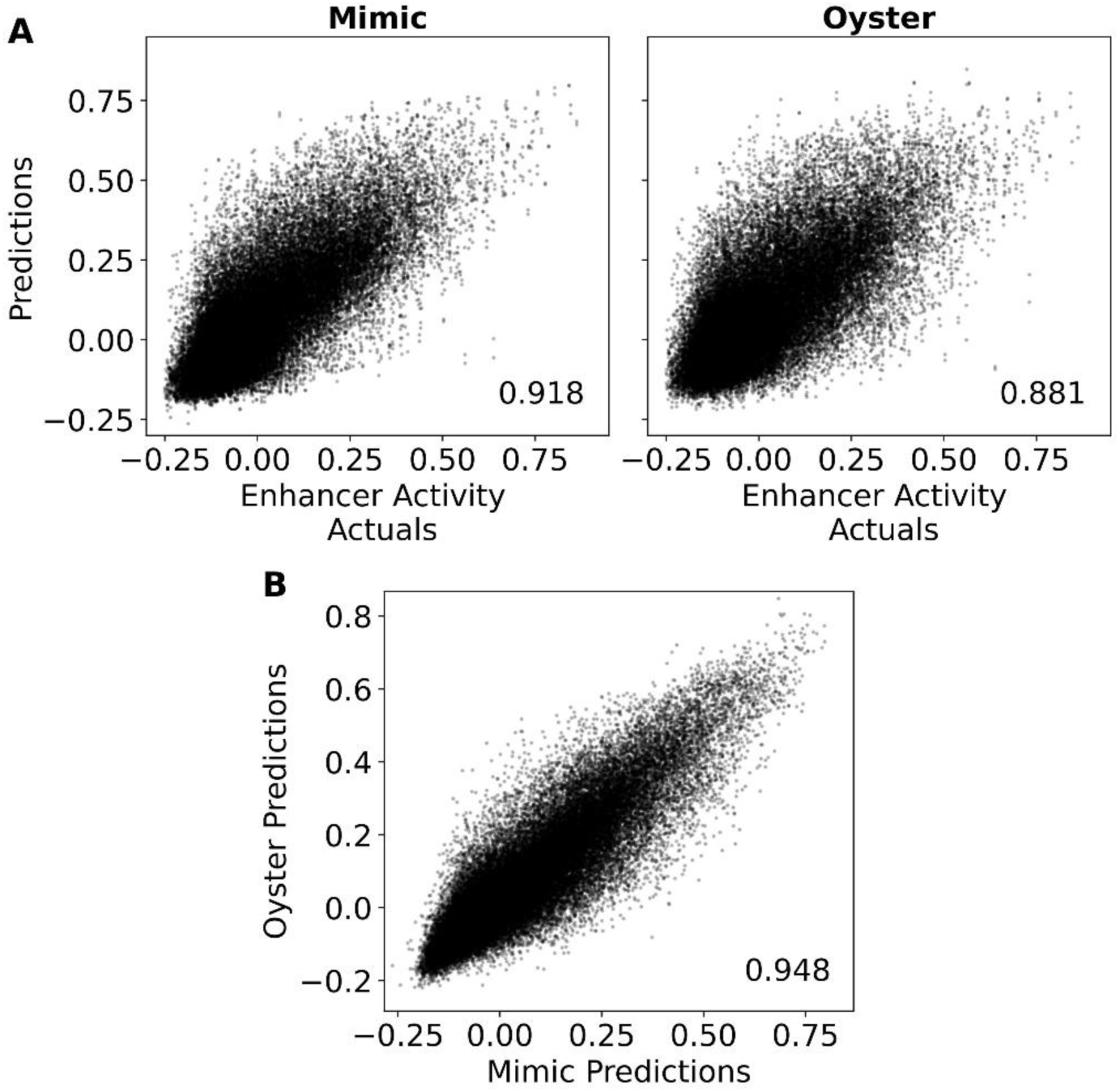
Mimic outperforms Oyster on modelling enhancer activity, but their predictions align strongly with each other. Scatter plots of weighted ensemble predictions of Mimic and Oyster on enhancer activity on the testing set compared to (**A**) actuals and (**B**) each other.

Of the roughly 400 human protein-DNA motifs analyzed, only a select few are strongly contributive, all of which contribute positively to enhancer activity (**Fig. 14**). The most contributive motifs are the ATF4 and CEBPG motifs which exhibit sequence similarity (**Fig. 15**, **Fig. 16**). This finding is consistent with the essential role of the Integrated Stress Response (ISR) in tumor survival where ATF4 and CEBPG are crucial effectors. ATF4 serves as the master regulator of the ISR (Harding et al., 2003) and forms heterodimers with CEBPG to facilitate stable DNA binding and gene regulation (Huggins et al., 2016). While the ISR is maintained at low basal levels in normal tissues, cancer cells often hijack this pathway to sustain proteostasis and metabolism within nutrient-deprived and hypoxic tumor microenvironments (Ye et al., 2010).

**Fig. 14.**
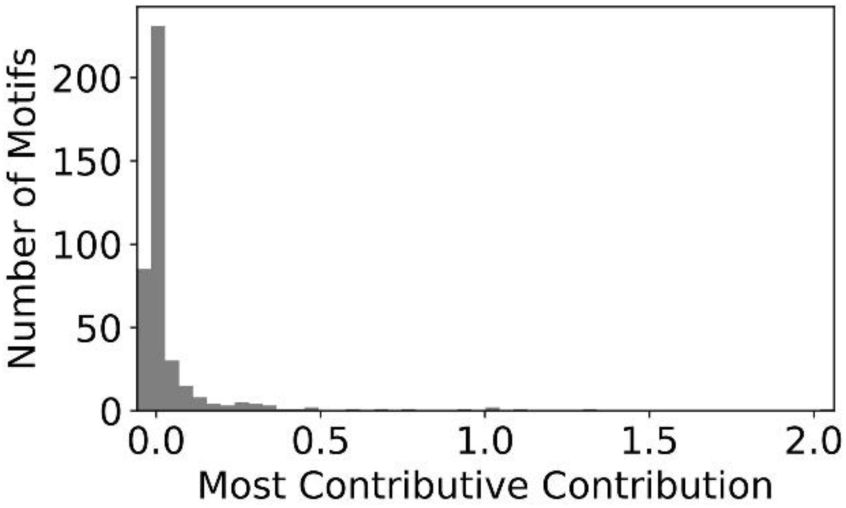
There are few strongly contributive motifs for modelling enhancer activity. See Fig. 9 for additional descriptions.

**Fig. 15.**
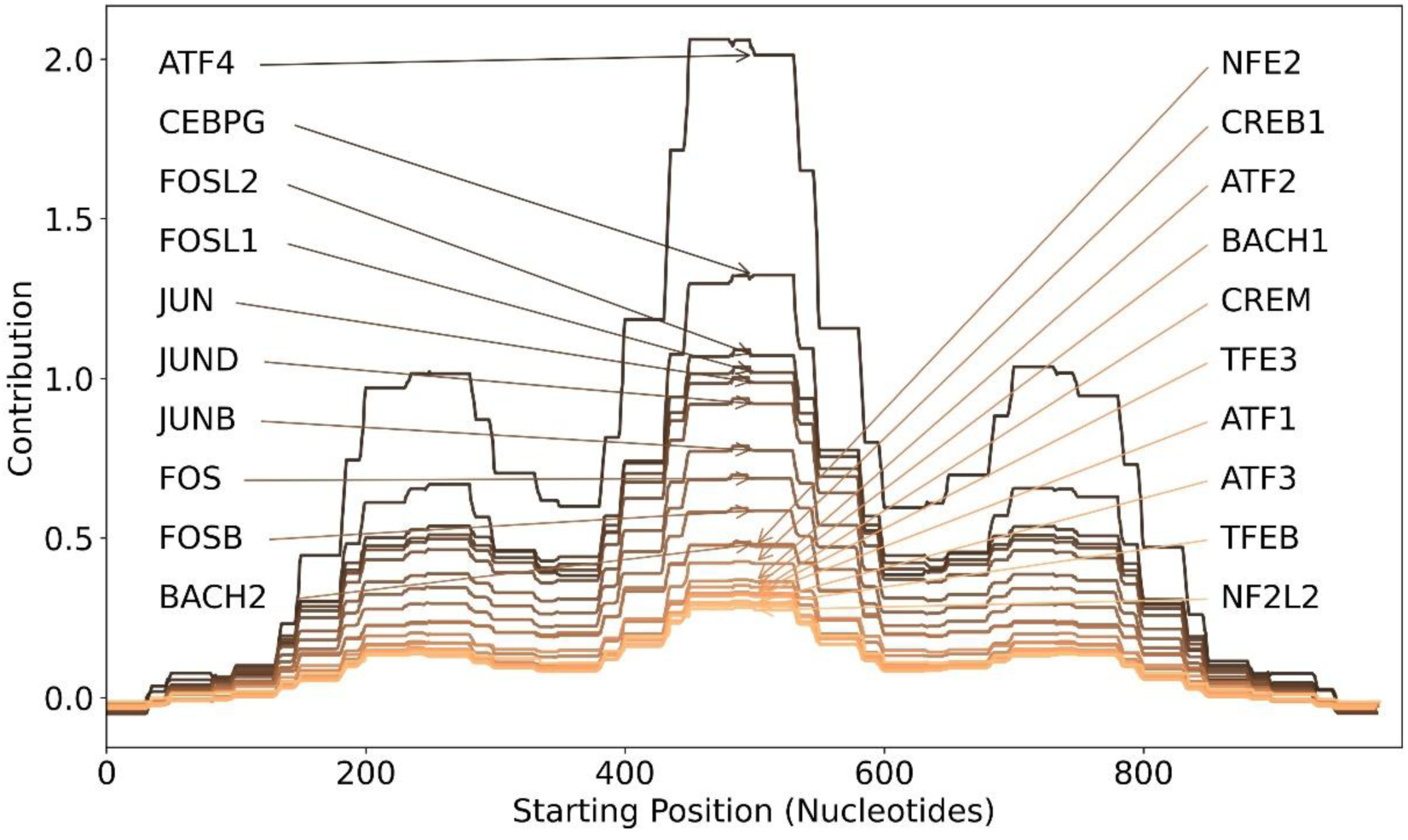
ATF4 is the most contributive motif for enhancer activity. Motif contributions of the top 20 most contributive motifs for each possible starting position of the 1000 nt sequence.

**Fig. 16.**
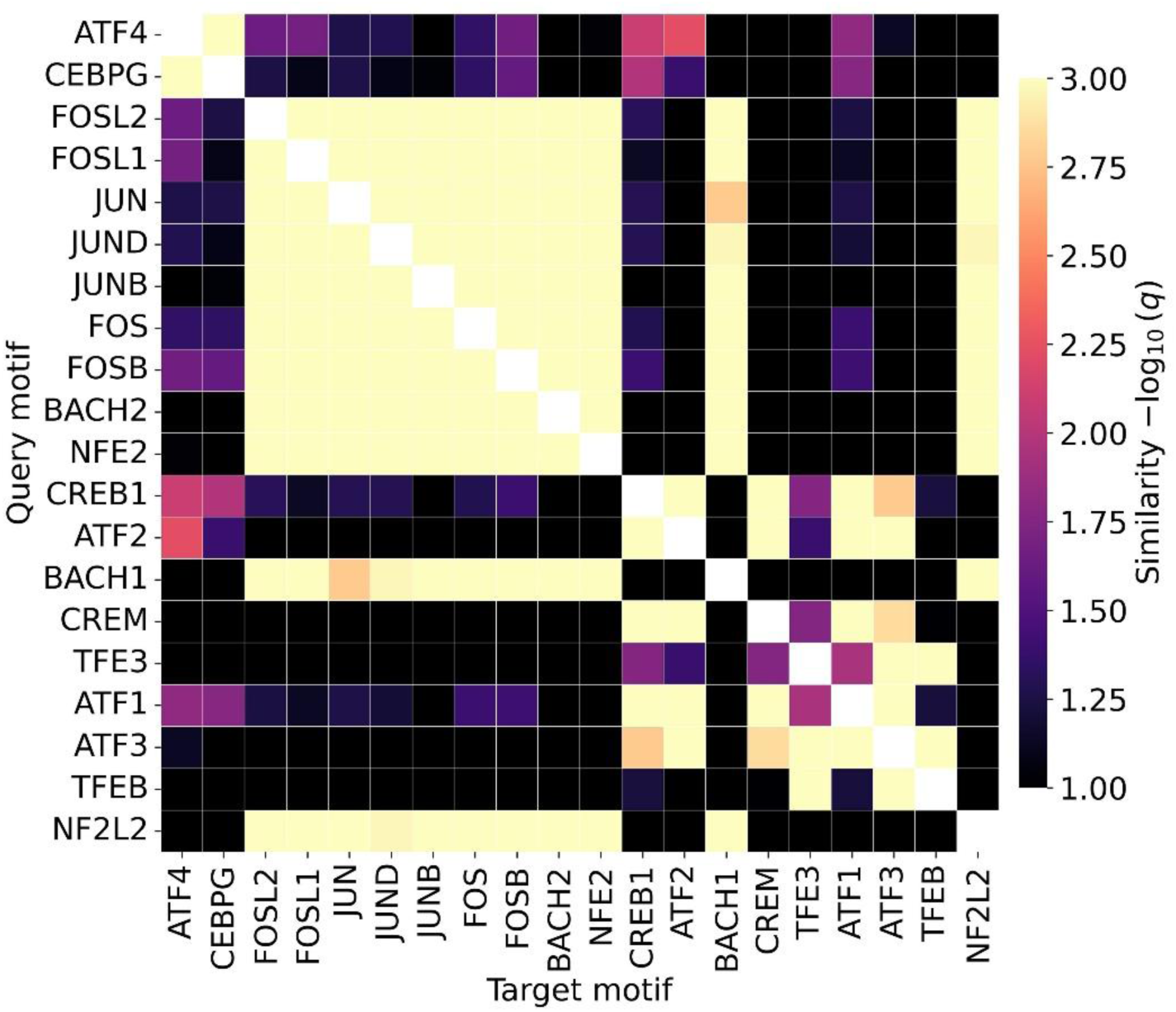
The most contributive motifs for predicting enhancer activity exhibit sequence similarity to one another. See Fig. 11 for additional descriptions.

The model also identifies the FOS and JUN motifs as strongly contributive motifs. FOS and JUN form the AP-1 (Activator Protein-1) complex, a pioneer transcription factor that open closed chromatin to allow gene regulation by other factors such as ATF4 (Biddie et al., 2011; Shaulian & Karin, 2002). NFE2 and BACH2 interact with similar DNA motifs which are involved with erythroid specific gene regulation (Andrews et al., 1993; Itoh-Nakadai et al., 2014), making them defining lineage factors for the K562 erythroleukemia cell line. Other ATF motifs—ATF2, ATF1, ATF3—are also strongly contributive. These proteins are involved in parallel stress-signaling and MAPK-mediated pathways, providing broader resilience to cancer cells (Hai & Curran, 1991).

Furthermore, the model identifies that contributions of all protein-DNA motifs have similar positional dependence. The contributions of the motifs are the greatest when present at the center of the sequence and generally decreases away toward the flanking regions. However, there are also two other considerable peaks at ±250 nts from the center. This is most likely an artifact from the data used for modelling. The site selection involved sliding windows and identifying windows with the greatest local score. Neighboring windows of ±250 nts and ±500 nts were added as translational augments. It is expected that the addition of more diverse neighboring windows would result in a unimodal positional dependence, a hypothesis that warrants future investigation.

Collectively, the Oyster model identifies an enhancer regulatory grammar that is fundamentally dependent on both the presence and spatial positioning of motifs critical for malignant survival and lineage identity in K562 cells.

## 5 General Discussion

Human understanding of genomics concepts is arguably limited to the contributions of *k*-mers and their positional dependencies. Oyster is a novel NN that learns *k*-mer composition and *k*-mer position but ignores *k*-mer associations, the first model to do so to the best of our knowledge. By comparing models that consider *k*-mer associations to Oyster, the impact of *k*-mer associations on genomics phenomena can be estimated. Furthermore, the position-specific contributions of any *k*-mer can be determined directly from model weights, making Oyster a glass-box model.

In case studies spanning two important genomics research topics of protein-DNA interactions and enhancer activity, Oyster models performed worse than Mimic models that considered *k*-mer associations. However, in both cases, Oyster’s predictions aligned strongly (recip-PCC > 0.9) with those of Mimic. This demonstrates the Oyster models utility as faithful proxies of the Mimic models and suggests that *k*-mer associations were weakly impactful to the specific phenomena investigated. To be clear, this is not to suggest that *k*-mer associations are negligible across all genomic phenomena including different protein-DNA interactions and enhancer activity tasks. Models that consider *k*-mer associations should be compared to Oyster models for all modelling tasks.

The Oyster models served as faithful proxies to Mimic models despite being trained on the data independently. An alternative approach is to train Oyster models using the Mimic model (or another model) predictions directly, essentially using Oyster as a surrogate model. It is expected that this would yield predictions that are even more aligned, but this remains to be tested.

As mentioned, *k*- and *z*-patterns can be generated by clustering *k*-mers by composition. An alternative is to cluster *k*-mers by their Oyster *k*-mer embedding, which projects *k*-mers based on both their composition and their relationship to the output. Such an approach could be used to refine existing DNA motifs and yield novel, biologically important *k*- and *z*-patterns. While this was beyond the scope of the current study, its potential for discovery warrants future exploration.

## 6 Acknowledgements

None.

## 7 Author Contributions

**Conceptualization:** Husam Abdulnabi

**Data curation:** Husam Abdulnabi

**Formal analysis:** Husam Abdulnabi

**Investigation:** Husam Abdulnabi

**Methodology:** Husam Abdulnabi

**Project administration:** Husam Abdulnabi

**Software:** Husam Abdulnabi

**Supervision:** J. Timothy Westwood

**Writing – original draft:** Husam Abdulnabi

**Writing – review & editing:** J. Timothy Westwood

## 9 Supplementary

**Sup. Text. 1.**
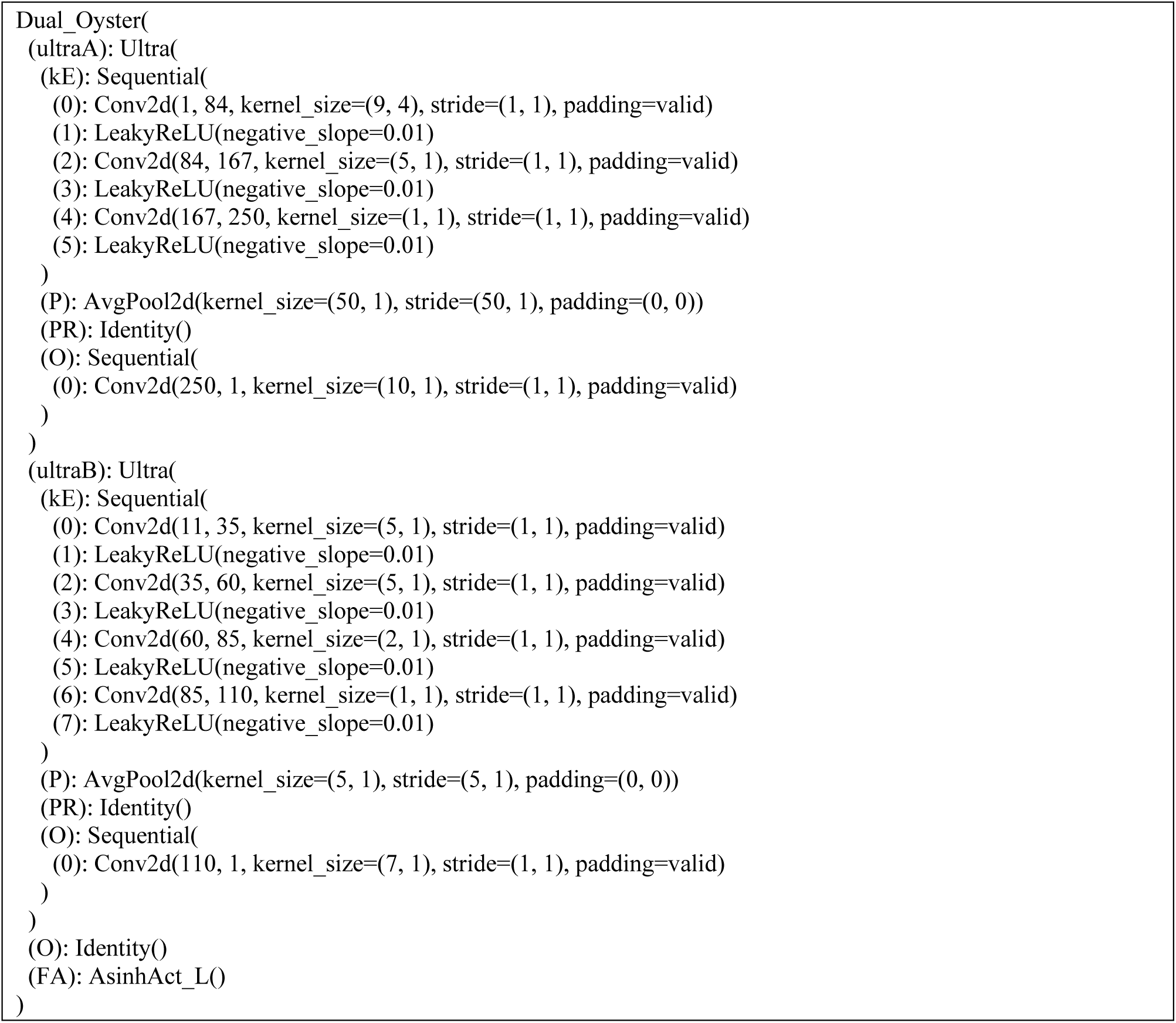
Optimized Multi-Oyster architecture for modelling CTCF-DNA interactions.

**Sup. Fig. 1.**
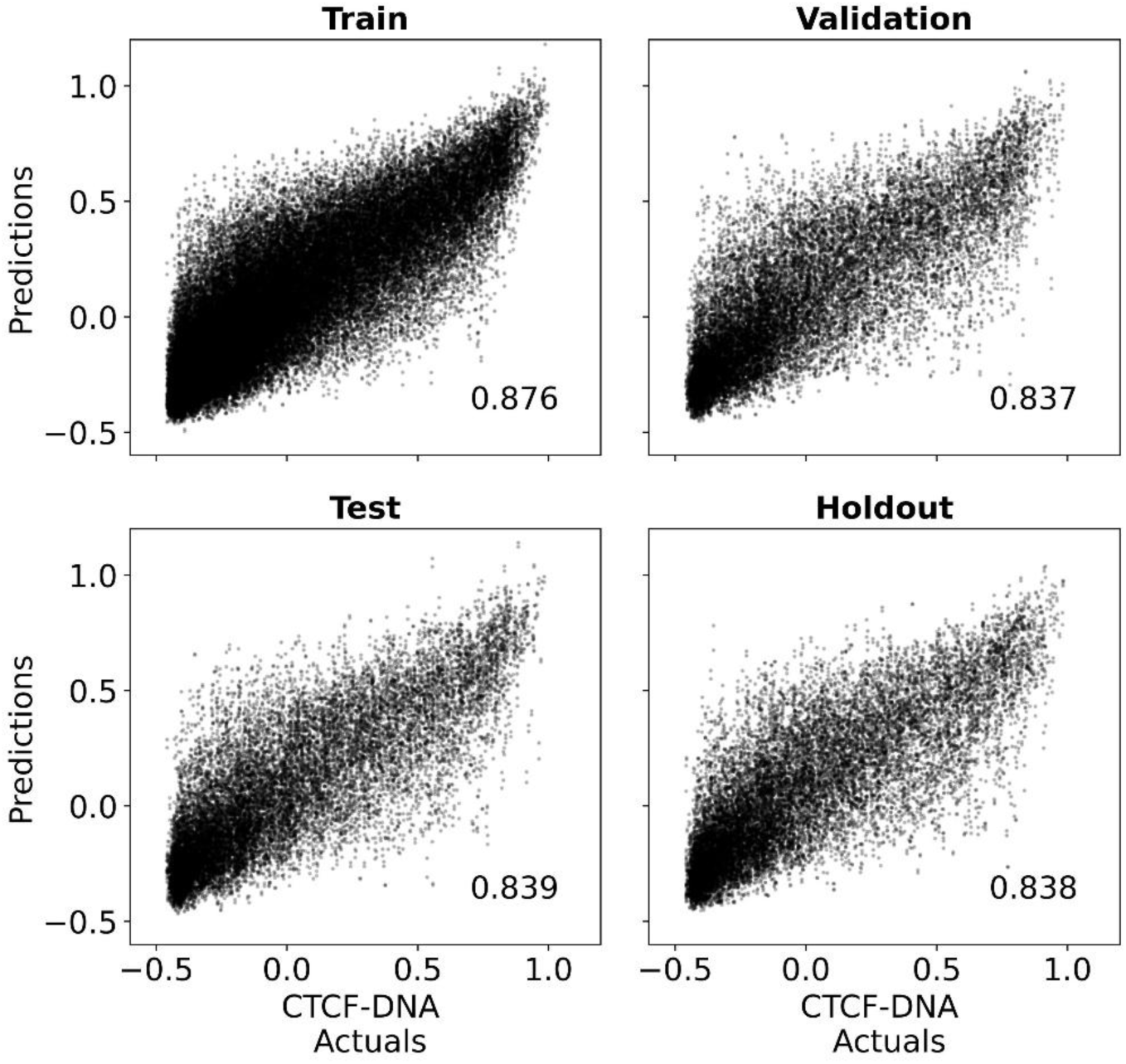
Multi-Oyster performs well on all CTCF-genome interactions sets. Scatter plots of weighted ensemble predictions of Multi-Oyster on CTCF-genome interactions on the training, validation, testing, and holdout tests compared to actuals. The recip-PCC is included at the bottom right of each plot.

**Sup. Text. 2.**
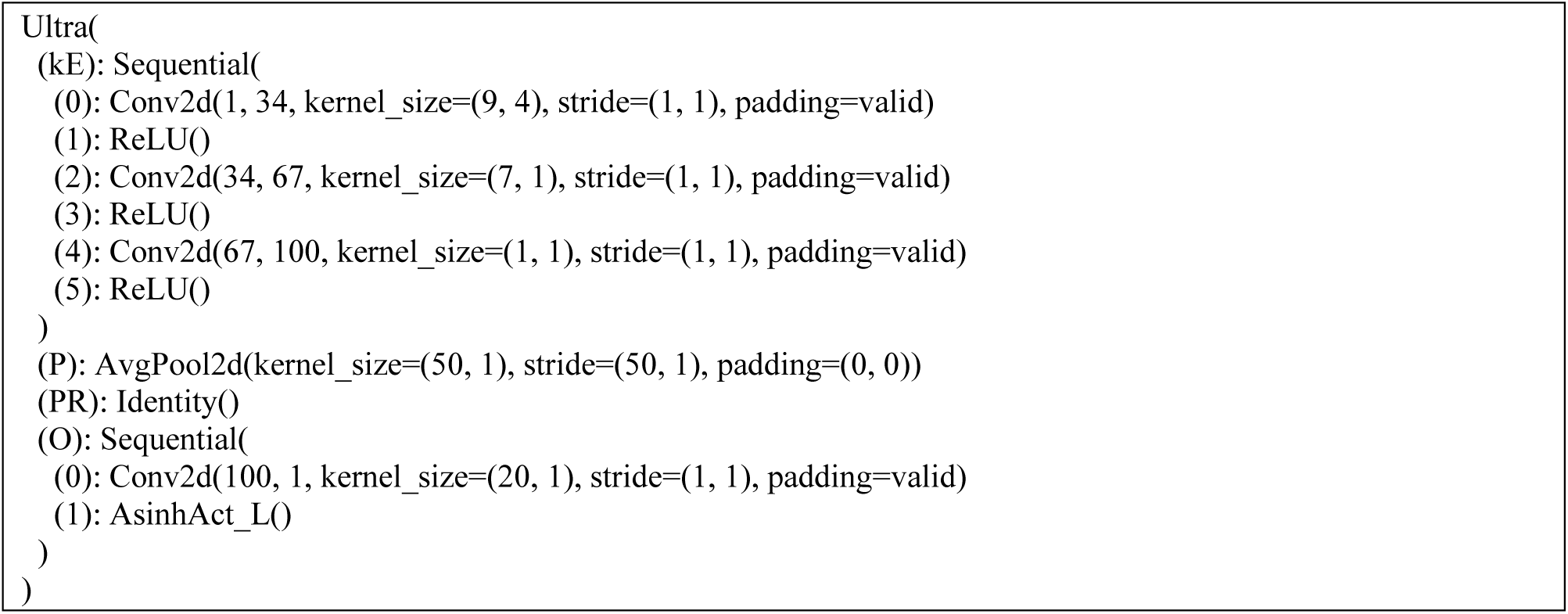
Optimized Oyster architecture for modelling enhancer activity.

**Sup. Fig. 2.**
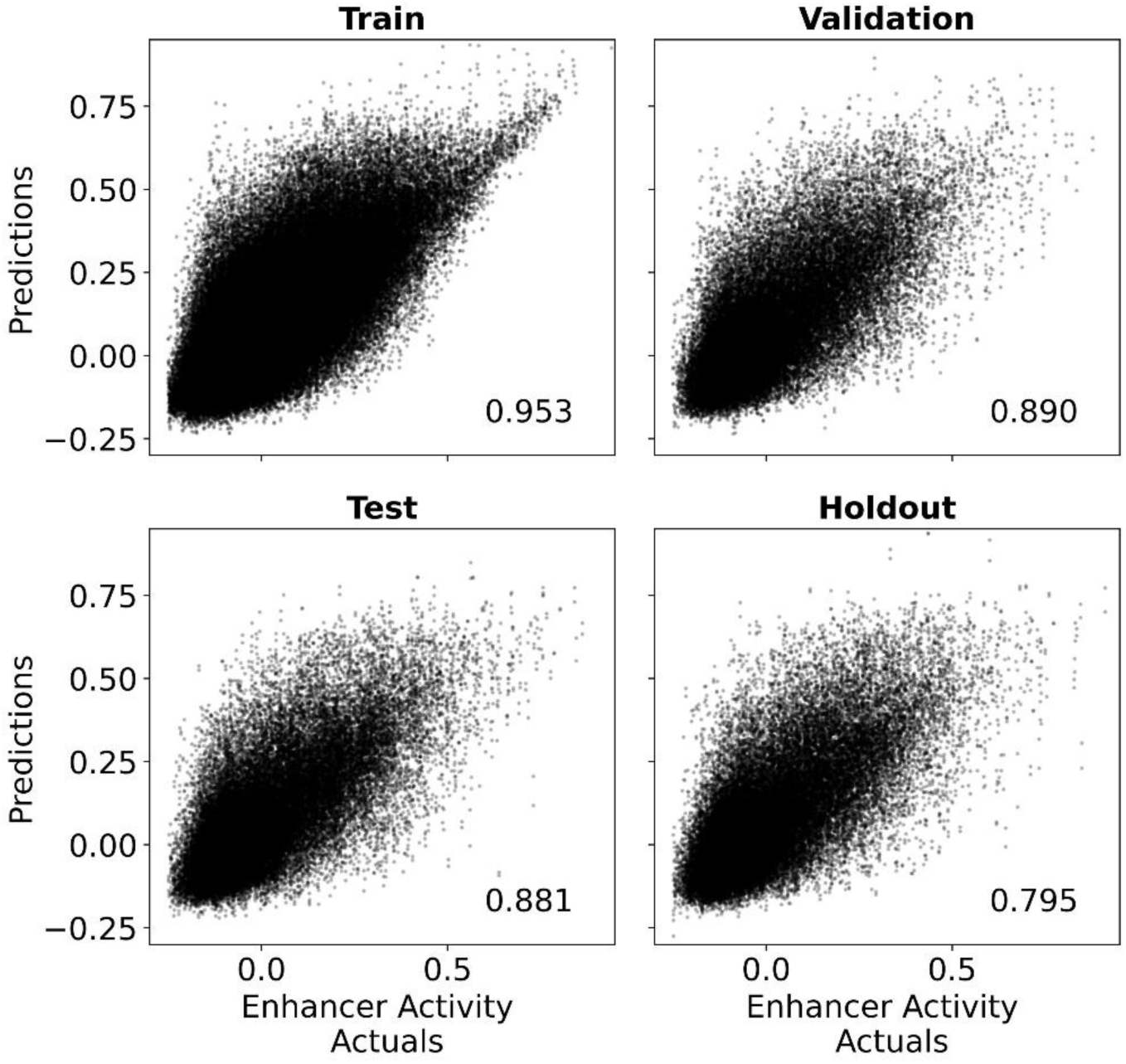
Oyster performs well on all enhancer activity sets. Scatter plots of weighted ensemble predictions of Oyster on enhancer activity on the training, validation, testing, and holdout tests compared to actuals. The recip-PCC is included at the bottom right of each plot.

## References

Abdulnabi, H., & Westwood, J. T. (2026a). Binning-based Methods for Model Fitting and Data Splitting Improve Machine Learning Imbalanced Regression. BioRkiv. 10.1101/2025.06.26.661884

Abdulnabi, H., & Westwood, J. T. (2026b). Mimic: A Unified Neural Network Architecture for Modelling Genomic Sequences and Module-level Comparison. BioRkiv. 10.64898/2026.10.26.761901

Ancona, M., Ceolini, E., Öztireli, C., & Gross, M. (2019). Gradient-Based Attribution Methods (pp. 169–191). 10.1007/978-3-030-28954-6_9

Andrews, N. C., Erdjument-Bromage, H., Davidson, M. B., Tempst, P., & Orkin, S. H. (1993). Erythroid transcription factor NF-E2 is a haematopoietic-specific basic–leucine zipper protein. Nature, 362(6422), 722–728. 10.1038/362722a0

Beer, M. A., & Tavazoie, S. (2004). Predicting Gene Expression from Sequence. Cell, 117(2), 185–198. 10.1016/S0092-8674(04)00304-6

Biddie, S. C., John, S., Sabo, P. J., Thurman, R. E., Johnson, T. A., Schiltz, R. L., Miranda, T. B., Sung, M.-H., Trump, S., Lightman, S. L., Vinson, C., Stamatoyannopoulos, J. A., & Hager, G. L. (2011). Transcription Factor AP1 Potentiates Chromatin Accessibility and Glucocorticoid Receptor Binding. Molecular Cell, 43(1), 145–155. 10.1016/j.molcel.2011.06.016

Chen, J., Zhang, Z., Li, L., Chen, B.-C., Revyakin, A., Hajj, B., Legant, W., Dahan, M., Lionnet, T., Betzig, E., Tjian, R., & Liu, Z. (2014). Single-molecule dynamics of enhanceosome assembly in embryonic stem cells. Cell, 156(6), 1274–1285. 10.1016/j.cell.2014.01.062

de Almeida, B. P., Reiter, F., Pagani, M., & Stark, A. (2022). DeepSTARR predicts enhancer activity from DNA sequence and enables the de novo design of synthetic enhancers. Nature Genetics, 54(5), 613–624. 10.1038/s41588-022-01048-5

ENCODE Project Consortium. (2012). An integrated encyclopedia of DNA elements in the human genome. Nature, 489(7414), 57–74. 10.1038/nature11247

Ghandi, M., Lee, D., Mohammad-Noori, M., & Beer, M. A. (2014). Enhanced regulatory sequence prediction using gapped k-mer features. PLoS Computational Biology, 10(7), e1003711. 10.1371/journal.pcbi.1003711

Guertin, M. J., & Lis, J. T. (2010). Chromatin landscape dictates HSF binding to target DNA elements. PLoS Genetics, 6(9), e1001114. 10.1371/journal.pgen.1001114

Hai, T., & Curran, T. (1991). Cross-family dimerization of transcription factors Fos/Jun and ATF/CREB alters DNA binding specificity. Proceedings of the National Academy of Sciences, 88(9), 3720–3724. 10.1073/pnas.88.9.3720

Harding, H. P., Zhang, Y., Zeng, H., Novoa, I., Lu, P. D., Calfon, M., Sadri, N., Yun, C., Popko, B., Paules, R., Stojdl, D. F., Bell, J. C., Hettmann, T., Leiden, J. M., & Ron, D. (2003). An Integrated Stress Response Regulates Amino Acid Metabolism and Resistance to Oxidative Stress. Molecular Cell, 11(3), 619–633. 10.1016/S1097-2765(03)00105-9

Huggins, C. J., Mayekar, M. K., Martin, N., Saylor, K. L., Gonit, M., Jailwala, P., Kasoji, M., Haines, D. C., Quiñones, O. A., & Johnson, P. F. (2016). C/EBPγ Is a Critical Regulator of Cellular Stress Response Networks through Heterodimerization with ATF4. Molecular and Cellular Biology, 36(5), 693–713. 10.1128/MCB.00911-15

Itoh-Nakadai, A., Hikota, R., Muto, A., Kometani, K., Watanabe-Matsui, M., Sato, Y., Kobayashi, M., Nakamura, A., Miura, Y., Yano, Y., Tashiro, S., Sun, J., Ikawa, T., Ochiai, K., Kurosaki, T., & Igarashi, K. (2014). The transcription repressors Bach2 and Bach1 promote B cell development by repressing the myeloid program. Nature Immunology, 15(12), 1171–1180. 10.1038/ni.3024

Jindal, G. A., & Farley, E. K. (2021). Enhancer grammar in development, evolution, and disease: dependencies and interplay. Developmental Cell, 56(5), 575–587. 10.1016/j.devcel.2021.02.016

Koo, P. K., Majdandzic, A., Ploenzke, M., Anand, P., & Paul, S. B. (2021). Global importance analysis: An interpretability method to quantify importance of genomic features in deep neural networks. PLoS Computational Biology, 17(5), e1008925. 10.1371/journal.pcbi.1008925

Kulakovskiy, I. V, Vorontsov, I. E., Yevshin, I. S., Sharipov, R. N., Fedorova, A. D., Rumynskiy, E. I., Medvedeva, Y. A., Magana-Mora, A., Bajic, V. B., Papatsenko, D. A., Kolpakov, F. A., & Makeev, V. J. (2018). HOCOMOCO: towards a complete collection of transcription factor binding models for human and mouse via large-scale ChIP-Seq analysis. Nucleic Acids Research, 46(D1), D252–D259. 10.1093/nar/gkx1106

Ng, P. (2017). dna2vec: Consistent vector representations of variable-length k-mers. ArXiv, arXiv:1701.06279.

Novakovsky, G., Fornes, O., Saraswat, M., Mostafavi, S., & Wasserman, W. W. (2023). ExplaiNN: interpretable and transparent neural networks for genomics. Genome Biology, 24(1), 154. 10.1186/s13059-023-02985-y

O’Geen, H., Squazzo, S. L., Iyengar, S., Blahnik, K., Rinn, J. L., Chang, H. Y., Green, R., & Farnham, P. J. (2007). Genome-Wide Analysis of KAP1 Binding Suggests Autoregulation of KRAB-ZNFs. PLoS Genetics, 3(6), e89. 10.1371/journal.pgen.0030089

Satopaa, V., Albrecht, J., Irwin, D., & Raghavan, B. (2011). Finding a“kneedle” in a haystack: Detecting knee points in system behavior. 31st International Conference on Distributed Computing Systems Workshops, 166–171.

Shaulian, E., & Karin, M. (2002). AP-1 as a regulator of cell life and death. Nature Cell Biology, 4(5), E131–E136. 10.1038/ncb0502-e131

Slattery, M., Zhou, T., Yang, L., Dantas Machado, A. C., Gordân, R., & Rohs, R. (2014). Absence of a simple code: how transcription factors read the genome. Trends in Biochemical Sciences, 39(9), 381–399. 10.1016/j.tibs.2014.07.002

Srivastava, D., Aydin, B., Mazzoni, E. O., & Mahony, S. (2021). An interpretable bimodal neural network characterizes the sequence and preexisting chromatin predictors of induced transcription factor binding. Genome Biology, 22(1), 20. 10.1186/s13059-020-02218-6

Wang, J., Zhuang, J., Iyer, S., Lin, X., Whitfield, T. W., Greven, M. C., Pierce, B. G., Dong, X., Kundaje, A., Cheng, Y., Rando, O. J., Birney, E., Myers, R. M., Noble, W. S., Snyder, M., & Weng, Z. (2012). Sequence features and chromatin structure around the genomic regions bound by 119 human transcription factors. Genome Research, 22(9), 1798–1812. 10.1101/gr.139105.112

Ye, J., Kumanova, M., Hart, L. S., Sloane, K., Zhang, H., De Panis, D. N., Bobrovnikova-Marjon, E., Diehl, J. A., Ron, D., & Koumenis, C. (2010). The GCN2-ATF4 pathway is critical for tumour cell survival and proliferation in response to nutrient deprivation. The EMBO Journal, 29(12), 2082–2096. 10.1038/emboj.2010.81

